# The importance of animal weapons and fighting style in animal contests

**DOI:** 10.1101/2020.08.26.268185

**Authors:** Alexandre V. Palaoro, Paulo Enrique Cardoso Peixoto

**Author notes:** Corresponding author: LUTA do Departamento de Ecologia e Biologia Evolutiva, Universidade Federal de São Paulo, São Paulo, Brazil. Rua Artur Riedel, 275, Eldorado, Diadema, CEP 09972-270.

## Abstract

In many species that fight over resources, individuals use specialized structures to gain mechanical advantage over their rivals during contests (i.e., weapons). Although weapons are widespread across animals, how they affect the probability of winning contests is still debated. According to theory, understanding the weapons’ function in contests depends on identifying differences in how weapons are measured (e.g., weapon length *versus* shape), and in how weapons are used during fights. Here, we developed a meta-analysis spanning 1,138 studies, from which were drawn 52 species and 107 effect sizes to identify: (1) what aspects of animal weapons are measured in the literature, and how these measures bias our knowledge; (2) how animals use their weapons during fights - i.e., weapon function; and (3) if weapon function correlates to the magnitude of how weapons influence contest resolution. First, we found that most of the literature focuses on linear measures of weapons, such as length. The few reports on weapon performance (e.g., biting force) were found only for Crustacea and Squamata. This bias highlights that measuring performance of weapons such as horns and spines might increase the breadth of our knowledge on weapons. Furthermore, we also found that linear measures showed stronger effects on contest success than performance measures. Second, we divided weapon function into displays and fighting style (i.e., how the weapon is used during fights). Regarding displays, most species displayed their weapons before contests (59.61%), rather than the body (34.61%). A minority (three species, 5.76%) engaged in fights without any type of display. Thus, species that bear weapons almost always perform displays before engaging in physical contact, a common hypothesis in contest theory that was never tested across taxa until now. Regarding fighting style, we found that most weapons were used for more than one behaviour during fights (e.g., squeezing and pushing). Further, pushing seems to be the most common behaviour among species, but it is usually accompanied by another behaviour, such as lifting or squeezing. Thus, oversimplifying fighting style can bias results because some styles might impose contrasting biomechanical pressures (e.g., pushing vs squeezing). Third, we found that display type did not influence the importance of weapon size on contests. Fighting style, on the other hand, influenced the effect of weapon size on contest outcome significantly. Species that used their weapons to impact, pierce or squeeze showed smaller differences between winners and losers when compared to pushing or lifting (and multifunctional weapons). Thus, pushing and lifting seem important for selecting larger weapons – even though some of them might also be used for squeezing, piercing or impacting. Overall, our results show that we have a biased understanding of animal weapons, built mostly on weapon size alone. Further, our analyses show that the importance of weapon size differs depending on the fighting style. If we lessen those biases, we will have a better and broader understanding of how weapons evolve and diversify.

## I. INTRODUCTION

Agonistic interactions have drawn attention from scientists since Darwin’s seminal publication of *The Descent of Man, and Selection in Relation to Sex* (Darwin, 1871). The field started mostly as descriptions of how animals engaged in agonism (e.g., Archer, 1988), but then gained a strong theoretical background by adding evolutionary game theory to its core (Maynard Smith & Price, 1973). Building on that theory, the field currently focus on how individuals make decisions during agonistic interactions (Hardy & Briffa, 2013; Chapin, Peixoto, & Briffa, 2019), the evolutionary consequences of winning or losing (Filice & Dukas, 2019), and what traits affect winning chances (Vieira & Peixoto, 2013). Among these diverse topics, however, one subject frequently raised is the existence of specialized morphological structures used during contests (termed animal weapons, McCullough, Miller, & Emlen, 2016). A weapon can be seen as specialized morphology used to gain mechanical advantage on the rivals (*sensu* Eberhard *et al*., 2018; see Table 1 for definitions), and because of this effect on the rival, weapons can affect all aspects of agonistic interactions. For instance, rivals may assess aspects of weapon size or performance when deciding whether to stay in or give up on the contest (Palaoro & Briffa, 2017; Pinto, Palaoro, & Peixoto, 2019). Weapons can also influence whether a contest escalate to physical fighting or not (Számadó, 2008). A myriad of studies have already shown that weapons are important for contest resolution, but most of them focus on single species (Vieira & Peixoto, 2013; Pinto *et al*., 2019). Reviews on this topic provide diverse insights on weapon evolution because they contain extensive knowledge on the shapes and sizes of animal weapons (Emlen, 2008; Rico-Guevara & Hurme, 2019). However, they lack quantitative information on selection pressures that may act on weapons. Due to this, it is necessary to integrate data of a wider diversity of species to estimate how weapons affect contests; hence providing a broader picture of the relative importance of weapons for contest resolution.

**Table 1.** Definitions used throughout this paper.

| Term | Definition |
| --- | --- |
| Weapon | Specialized morphology used to gain mechanical advantage on rivals during physical contact. |
| Display | Any behaviour or movement that can be used as a cue of fighting ability, or as a threat behaviour. In most species, it is the part of the contest that does not involve physical contact. In the species where rivals are in physical contact, it is associated with physical contact of low intensity, such as antennae touching. |
| Fight | Part of the contest where the individuals are within reach of one another and are in physical contact. Can be seen as a physical struggle between two rivals. |
| Contest | Aggressive competition between two rivals of the same species for an indivisible resource. Can be composed of two phases: a display phase and a physical contact phase. |
| Fighting style | Behaviours performed during physical contact that aim to gain mechanical advantage over the rival. Can be composed of either a single or multiple behaviour used simultaneously. Frequently associated with movements of the weapon itself (e.g., squeezing). |
| Weapon function | The combination of display and fighting style an individual uses during a contest. See Table 3 and Table S1 for a detailed list of weapon function. |

One major hurdle to assess the relative importance of weapons is the large diversity of measures used as proxies of weapon traits (Vieira & Peixoto, 2013). For instance, weapon traits may be divided into three major categories: performance (such as a weapon’s capacity to exert force), size (such as weapon mass or length), and shape (such as the ratio between weapon length and width). Although all of them may be important for determining the winner, there is a debate on which trait(s) should influence contests the most (Eberhard *et al*., 2018; Palaoro *et al*., 2020). If one trait is more important than the others, we can expect stronger selective pressure on that trait. However, a systematic review on how different weapon traits affect winning chances is still lacking. A first step would be to identify the existing patterns on how studies are measuring the traits of different weapons to assess if there are any gaps or potential biases in our knowledge. The second step would be to quantify the relative importance of each weapon trait for contest resolution. For example, if performance is more important than size for victory, we expect larger differences between winners and losers on proxies of weapon performance (such as weapon muscle mass) when compared to proxies of weapon size (such as weapon length).

Another point frequently raised in studies of weapon evolution is related to how the function of the weapon should influence the importance of each weapon trait on winning contests (McCullough, Tobalske, & Emlen, 2014; McCullough *et al*., 2016, see Table 1 for definitions). Before physical contact, for instance, individuals may vary on whether they use their weapons as displays or not. Rivals can either use their weapons before the fight as threat displays (Lappin *et al*., 2006), or use their weapons only during physical contact (Katsuki *et al*., 2014). Weapons also vary in how they are used during fights, i.e., fighting style (*sensu* McCullough *et al*., 2016). For instance, individuals may use weapons to push (Graham, Padilla-Perez, & Angilletta, 2020), squeeze (Dennenmoser & Christy, 2013), pierce (Candaten *et al*.,2020), or even lift and throw their rivals away from the contested resource (Goyens, Dirckx, & Aerts, 2015). Due to this variation, fighting styles have been notoriously difficult to assess precisely. However, without a standardized framework to assess weapon function, we cannot identify potential differences in the selective pressures on weapon traits that may be related to the way rivals use their weapons during contests. For this, it is also necessary to perform a systematic review of how individuals use their weapons during contests; then test whether different fighting styles are able to predict how important each weapon trait is for contest resolution.

A crucial step to perform a systematic review to estimate the importance of weapons in contest resolution is to quantify how weapon traits affect winning chances. One way to obtain such quantification is to estimate how much winners and losers differ in a given weapon trait (e.g., weapon size or performance). However, since different species may differ in how much a given weapon trait varies or in the type of measure that can be done, the estimation of the relative difference in weapon traits among winners and losers must be standardized to be comparable. In particular, the standardized mean difference in weapon traits between winners and losers is a powerful metric frequently used in ecology and evolution (i.e., Hedges’ *g* effect size, Nakagawa & Cuthill, 2007): larger standardized mean differences between the groups indicate a stronger effect of the measured trait or variable. When applied to animal contests, it would be expected that traits that are more important in resolving contests should show marked differences between winners and losers, while traits that are less important should be more similar between rivals (e.g., Vieira & Peixoto, 2013). Therefore, as specific weapon traits become more important to determine contest resolution, the greater should be the difference in such traits between winners and losers.

Here, our goal is to quantify the importance of weapon traits and fighting style on contest resolution. For this, we performed a meta-analytical review to answer the following questions: 1) Are there biases in how weapons are measured and do distinct traits differentially influence contest resolution? 2) How much variation exists in weapon function and are there similarities among species? 3) Do similarities in displays and fighting style influence the relative importance of weapon traits in contests? Below we provide a general description of how we gathered information to answer these questions and then present our rationale and findings for each question separately.

## II. GENERAL PROCEDURES

Below we describe how we searched for the articles used in this review, how we extracted and transformed the information obtained from the articles and how we controlled for phylogenetic effects in each analysis. Such procedures were the same to obtain the data used to answer our three questions. Specific procedures adopted for each question are separately described in the corresponding section.

### (1) Study selection and data gathering

We searched for articles using the Web of Science (https://www.webofknowledge.com) and Scopus (https://scopus.com) using their core collection databases from 1945 to 2019. For both searches we used the following keywords: “contest*”, “fight*”, “assessment*”, “resource holding p*”, “resource-holding-p*”, “agonis*”, “territory defen*e”, “weapon*”, “armament*”, “sexual* trait*”, “sexual*-selected trait*” “body size*”, “antler*”, “horn*”, “jaw*”, “claw*”. All keywords were used with the “OR” Boolean operator. During our search, we excluded all studies in which the species did not bear a weapon (Table 1) such as butterflies and dragonflies. Since we had to classify species according to how individuals use them during contests, we excluded all species in which the behaviours adopted during the contests were undescribed.

For the selected studies, we collected information about mean values (and their corresponding variation) of weapon measures for winners and losers of contests. Within each study, we also recorded the pairing method used by researchers. We distinguished between studies in which fighting individuals were experimentally paired to have similar body sizes but differing weapon sizes and studies in which individuals were randomly chosen to contest (we also included in this second group studies in which one individual was free to choose their rivals). Further, we found no studies that paired individuals by weapon size and let body sizes differ; thus, we use ‘paired contests’ to refer to contests where individuals have similar body sizes, but differing weapon sizes. Another confounding effect might come from resource value (*sensu* Arnott & Elwood, 2008): individuals that value more a resource often are more motivated to fight regardless of their fighting ability and have a higher chance of winning (e.g., Palaoro *et al*., 2017). Therefore, to avoid any bias related to resource value, we only included the treatments in which there was no evidence that individuals had different motivations to fight. We also recorded whether the study was performed in a laboratory environment or in the wild.

We obtained a total of 1108 papers through those searches. We also added 30 relevant papers cited in reviews that we did not find in our primary searches (Emlen, 2008; Vieira & Peixoto, 2013; Pinto *et al*., 2019; Rico-Guevara & Hurme, 2019). After excluding papers that did not provide all necessary information (Fig. S1), our final data set comprise 48 papers that contained 52 species, comprising both vertebrate and invertebrate species (Fig. S2). Within these, we had information for 33 species involving randomly paired rivals and 27 species involving rivals paired by size (Fig. 1). We performed all steps of the literature review following the PRISMA protocol (Liberati *et al*., 2009), and the flow diagram containing all steps can be found in Fig. S1.

**Fig. 1.**
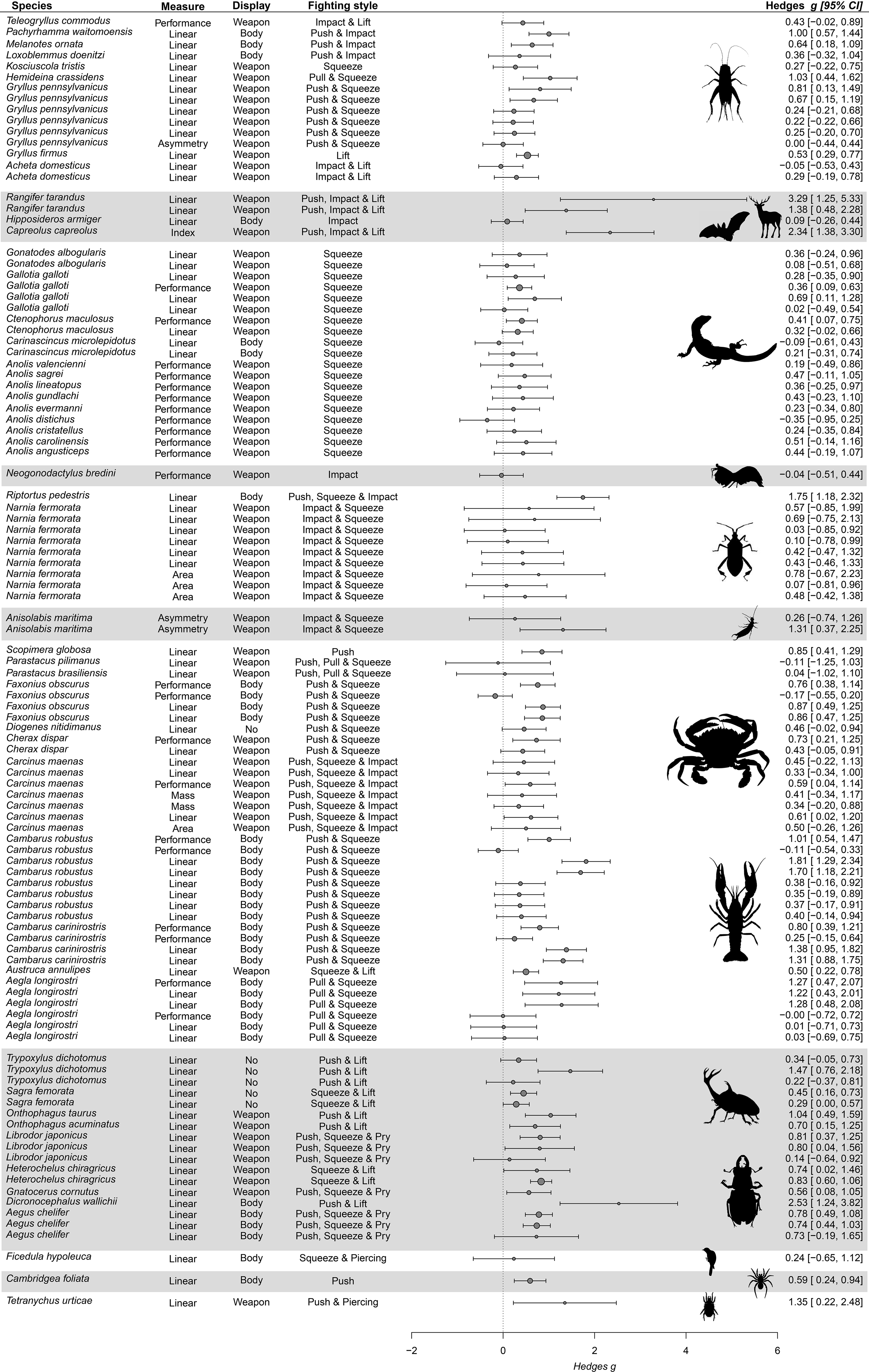
Hedges’ *g* value (circle) and corresponding confidence intervals (lines) for species, pairing method (whether contests were paired for body size or not), component measured (e.g., length, width, mass), type of display (i.e., displaying weapon, body size, or not displaying) and function of the weapon during the contests (see topic 3). Study demonstrates which samples were taken from the same study (references in Supplementary File S1). Larger grey circles denote studies with larger sample sizes. Silhouettes were taken from phylopic. Grey rows were selected randomly to facilitate visual separation of groups. For more information regarding the studies and pairing methods, see Fig. S3.

### (2) Effect size calculations and phylogenetic tree

To compare the magnitude of trait differences between winners and losers across different species, we used Hedges’ *g*, which is a standardized dimensionless measure that allows comparisons among different types of measurements and species (Koricheva, Gurevitch, & Mengersen, 2013). To calculate Hedges’ *g*, we used the mean trait values, their standard deviations, and respective sample sizes for each trait measure (such as weapon mass or weapon length) of winners and losers of each selected study. Since we always calculated the Hedges’ *g* as the difference between winner and loser traits, values greater than zero indicate that winners, on average, have a greater trait value than losers; while values smaller than zero indicate that losers have, on average, a lower trait value than winners. Unfortunately, mean values and standard deviations were not always available on papers. In those cases, we gathered results from statistical tests comparing winners and losers (e.g., *t*-values, F-values, degrees of freedom, and sample sizes) to transform the statistical results into Hedges‘ *g* values using the package “*compute.es*” in R (Del-Re, 2013). Whenever the mean and standard deviations were displayed on graphs, we used the *webplotdigitizer* software to extract the values directly from the figures (Rohatgi, 2019). If none of those options were available, we contacted the corresponding author to request the data.

To control for the phylogenetic relatedness between the species in our sample, we built a phylogeny comprising all 52 species (Fig. S2) using the Interactive Tree of Life online tree generator (iTOL; https://itol.embl.de/). After generating the tree, we estimated branch length using a Brownian motion model of evolution to simulate an ultrametric phylogenetic tree (Paradis, Claude, & Strimmer, 2004). We then transformed the ultrametric tree into a variance-covariance matrix that reflects the phylogenetic relatedness among the species. The variance-covariance matrix was then imputed in our meta-analytical models (see below for a detailed description of each model) as a random variable. We made these procedures using the packages “*rotl”* (Michonneau, Brown, & Winter, 2016) and “*ape*” in R (Paradis *et al*., 2004).

## III. WHAT IS THE EFFECT OF DIFFERENT WEAPON TRAITS ON CONTEST RESOLUTION?

Weapons can be measured regarding their performance, size, and shape; each of these traits can be used as proxies for fighting ability (see Vieira & Peixoto, 2013 and Palaoro *et al*., 2020b). This diversity of measures sparked interest on whether (and how) distinct weapon traits may affect contest resolution (Lappin & Husak, 2005; McCullough *et al*., 2016; Eberhard *et al*., 2018; Palaoro *et al*., 2020). Given the important role weapons play in deciding a contest winner, understanding if each trait influences contest resolution is thus an important step to reveal how selective pressures may have acted on weapon evolution. In particular, if specific weapon traits are more important in determining victory, they should show greater differences between winners and losers when compared to weapon traits that are less important. For this reason, we used a meta-analytic review to assess how weapon traits are measured in studies on contest resolution; and tested if the difference between winners and losers changed according to the type of measurement made.

We divided the traits measured in six categories: weapon asymmetry, index of weapon size, weapon area, weapon performance, weapon linear measures (length or width), and weapon mass. Weapon asymmetry is often measured as the difference in size between the two sides of a bilateral weapon, e.g., the difference in the maxillae of a cricket (Briffa, 2008). The index of weapon size is used to calculate the size of a complex shape, i.e., to incorporate the complexity of branching of the antlers of a deer species into a metric of overall size (Hoem *et al*., 2007). Within performance, we considered only measures related to force output, such as muscle size and bite force measurements. Regarding area, linear measures, and mass, although they might represent the same component of a weapon (i.e., size), they have different scaling properties that can add non-random biases to the estimates (see Houle *et al*., 2011; Pélabon *et al*., 2014). In addition, in Arthropods, while mass may vary with individual condition, size is a fixed attribute in adults (e.g., Peixoto & Benson, 2008). Therefore, we separated linear from area and mass measurements.

### (1) Methods

After determining the type of trait measured for each species, we built a multilevel meta-analytical model using the type of trait as our moderator, the Hedges’ *g* effect size as our response variable, and the inverse of the variance of Hedges’ *g* as a weight. We also included five variables as random effects. First, we used ‘study ID’ because sometimes we extracted more than one effect size per study. In this random effect, we frequently had several traits of the same weapon being measured (e.g., linear and mass measurements within the same study). Thus, we build a correlation matrix for the ‘study ID’ random effect because the correlation between effect sizes can bias the outcome (e.g., Weaver *et al*., 2018; Mathot, Dingemanse, & Nakagawa, 2019). The correlation matrix had the Pearson’s correlation coefficient between different weapons traits to control for any allometric effect on our estimates. We found most of the correlations in the same papers from which we found the effect sizes. But, for those that we did not find, we searched the literature and used Pearson’s coefficients in the papers cited in Table S1. In the few cases in which no information was available, we used 0.5 as a coefficient value (following Weaver *et al*., 2018). The matrix we used can be assessed together with our codes and dataset (check the Data Availability section). Second, we used ‘species ID‘ to account for effect sizes that came from the same species, but different studies. Third, we used the environment in which the original study was performed (i.e., laboratory or wild). Fourth, we used a matrix containing the phylogenetic relatedness among species. Finally, we used the pairing system used in the original study (i.e., whether individuals were paired according to their body size or not).

To estimate heterogeneity and biases in the model, we used two approaches. First, we calculated the ratio of heterogeneity to the total variation observed across effect estimates in multivariate studies (I², Borenstein, 2009). We also partitioned the I² into the contribution of each random variable (Nakagawa & Santos, 2012). Thus, we had estimations of the within-studies variance (I²_study_, similar to most mixed models), the species ID variance (I²_species_), the phylogenetic variance (I²_phylogeny_), and the pairing method variance (I²_pairing_). The sum of these different I² are equal to the total variance observed (I²_total_). To estimate the phylogenetic signal in the effect size, we also calculated the phylogenetic heritability index, *H²*, which is similar to Pagel’s λ (Nakagawa & Santos, 2012). Finally, to test for publication bias, we conducted a modified version of the Egger’s test, in which we used the residuals of our meta-analytic model as the response variable and the standard deviation of the effect sizes as our predictor variable (Egger *et al*., 1997). If the intercept of this regression is not different from zero (α > 0.10), then there is little evidence for publication bias (Nakagawa & Santos, 2012).

### (2) Results

Linear measures were the most common trait found in the literature (74 out of 107 effect sizes, 69.15%). Performance measures were the second most common with 23 effect sizes (21.49%), followed by area (n = 4, 3.73%), asymmetry (n = 3, 2.88%), mass (n = 2, 1.87%), and index measures (n = 1, 0.9%). Linear measures were found for most species in the sample, but performance measures were concentrated on crustaceans and lizards. Of the 23 effect sizes on performance, 11 came from crustaceans (47.8%), 11 from lizards (47.8%) and 1 from a cricket (*Teleogryllus commodus*, Fig. 1). Therefore, there is a clear bias on the type of inference drawn for most groups: inferences still rely mostly on size measures, rather than performance, or other measures.

The low sample sizes for measures of weapon area, asymmetry, mass, and index would render any statistical assessment of their influence on contest success weak. Thus, we removed these measures from our sample and only tested the linear and performance measures. Overall, winners had greater weapon traits than losers regardless of whether linear or performance was measured (QM_2_ = 117.05, p <0.0001). However, linear measures had a greater effect on contest success than performance measures (QM_1_ = 23.29, p <0.0001; Fig. 2). The model showed low heterogeneity. Study ID and pairing method were responsible for most of that heterogeneity, while phylogeny and genus had negligible effects on variance (Table 2). We found evidence for publication bias on the effect sizes (Egger’s test; intercept: -0.315, 95% CI: -0.569, -0.06, t = -2.461, p = 0.015).

**Fig 2.**
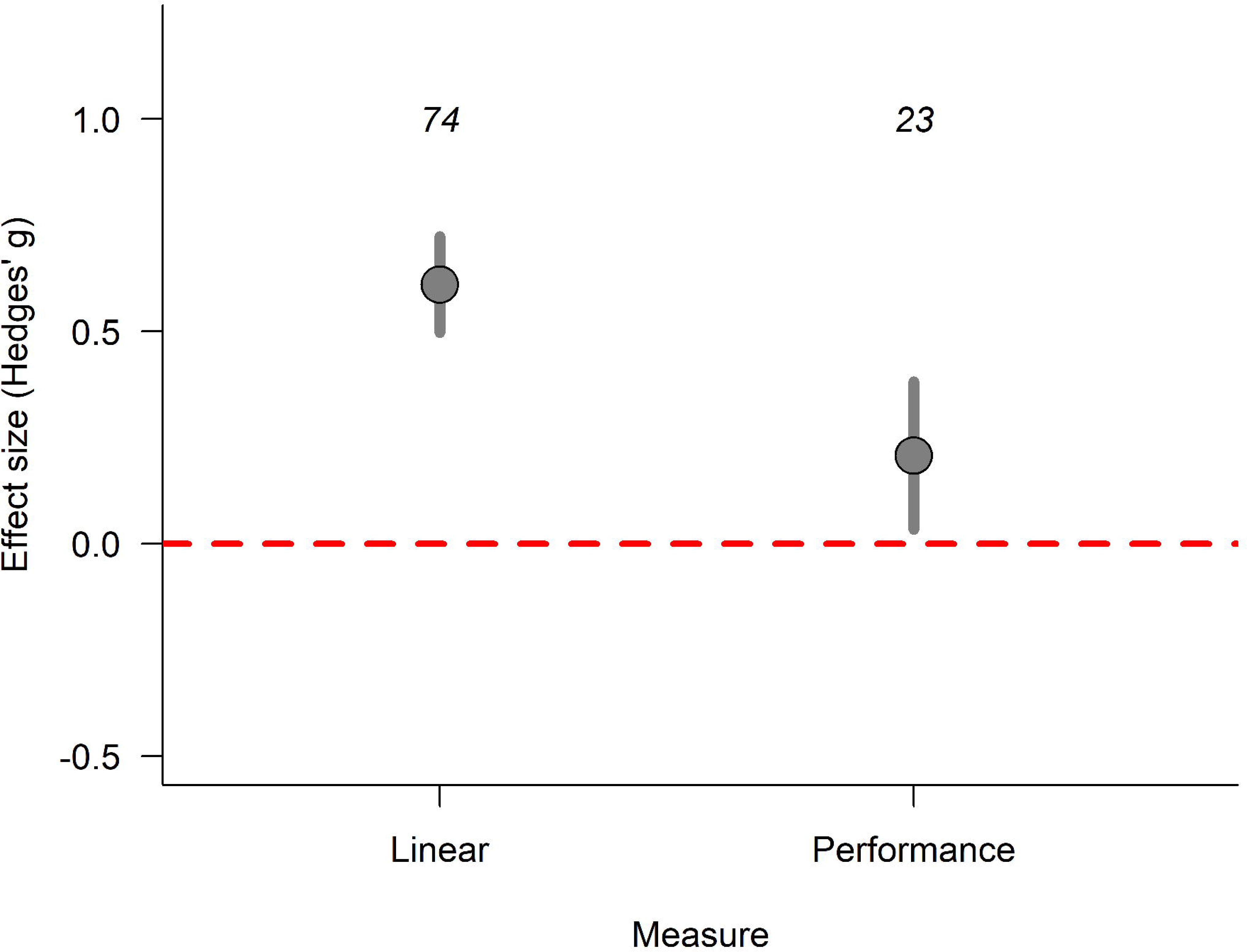
Weapons had an overall positive effect on contest success with winners having greater trait values than losers. Linear measures had a higher difference between winners and losers than performance measures demonstrating that winners tend to have much larger, rather than stronger weapons, than losers (QM_1_ = 23.29, p < 0.001). The effect size, Hedges’ g, represents the mean standardized difference between winners and losers. Positive values denote that winners were larger than losers, while negative values represent the opposite. Dots represent the estimated values from a multilevel meta-analytic model considering Hedges’ *g* as the response variable, weapon component as a moderator variable, and study ID, species ID, phylogeny, environment and pairing method as random variables. The error bars represent the 95% confidence interval of the estimate. The numbers above the error bars represent the number of effect sizes in each component.

**Table 2.** Heterogeneity of the multilevel meta-analytical model of differences in how weapons are measured.

| Random effect | I<sup>2</sup> (95% CI) |
| --- | --- |
| Study ID | 10.93 (9.009, 12.85) |
| Pairing method | 17.37 (15.44, 19.29) |
| Phylogeny | 0.012 (0.00, 1.92) |
| Genus | 0.0001 (0.00, 1.92) |
| Total | 23.71 (21.79, 25.63) |
| *H*<sup>2</sup> | <0.001 (0.00, 1.92) |

### (3) Discussion

Our results suggest that linear measures have a higher relative importance to determine contest outcome than performance measures. Therefore, different traits of the weapon may provide distinct information about the relative importance of weapons in contests. Given recent evidence (e.g., Palaoro *et al*., 2020b; Emberts *et al*., 2021), it is somewhat surprising to find that weapon size is more important than weapon performance on contest resolution. However, interpreting these results requires caution because of the biases and limitations of the current literature. For instance, performance measures are concentrated on crustaceans and lizards (only one performance measure was found outside those two groups). These measures provide information on the performance of weapons that have very similar morphologies: claws and jaws. Meanwhile, linear measures have a wider diversity of weapons, such as antlers, mandibles, and horns, which are also distributed among more taxa. Therefore, to ensure that the effect we found is not an artifact associated with differences in the diversity of weapons that have linear and performance measures, we reran the analysis using only data gathered from crustaceans and lizards. The result was the same to that observed in the analysis considering all species: the effect size of linear components was higher than the effect size of performance components (Fig. S4). Thus, despite the bias in the diversity of weapons, it seems that linear measures are more important for contest resolution than performance measures.

One explanation for why linear measures seem more important than performance measures may reside on engaging the rival from a safe distance (Eberhard *et al*., 2018). Being able to handle the rival without being exposed can also give the individual more time align the weapon relative to the rival’s weapon or body without incurring any extra costs (fighting skill, Briffa & Lane, 2017). If that is true, then any weapon can benefit from being larger. However, that may be particularly true for weapons that can push the rival (even if pushing co-occurs with other behaviours, see the next session). That way, the individual that has more time to find a better spot to lock on the rival and enjoy and increased chance to win the contest by pushing the opponent better. Thus, individuals might benefit from having a larger weapon, regardless of their weapon morphology. Alternatively, weapons might be used as visual displays and during fights to gain mechanical advantage on the rival. These selective pressures combined could favour bigger weapons if size influences contest success by getting rivals to retreat before physical contact (McCullough *et al*., 2016). We tackle this hypothesis in the next sessions.

Along with the lack of measurements of performance during fights, there is a clear information gap on the performance of weapons that are used for other types of behaviours, such as striking or ramming. The only example outside of biting performance we have on our dataset are the bullet-fast strikes of mantis shrimps (Green & Patek, 2015). Most of the types of weapons where we miss information on other measures are, in fact, weapons that do not have specific muscles attached to them, such as antlers, horns, and spines. These weapons are used for a multitude of behaviours during fights (see next session) and rather than having their own movement (i.e., biting, striking), they are used simultaneously with body movement. A few studies show how to measure the performance of horns (McCullough, 2014) and spines (Crofts *et al*., 2019) during relevant tasks. Thus, expanding our knowledge to how much these performances influence contests can broaden our understanding of weapon evolution.

## IV. ARE THERE SIMILARITIES AMONG SPECIES IN WEAPON FUNCTION?

Weapon function may be composed of two moments. In the first moment (not present in some species, see below), individuals may use their weapons as displays before physical contact ensue. In the second moment, individuals use their weapons to manipulate the rival, which is referred here as fighting style (*sensu* McCullough *et al*., 2016). Both moments can vary broadly within and between species. For instance, in the fiddler crab, *Austruca mjoebergi*, males use their claws as threat displays before physical contact. If the contest escalates, males use their claws to pinch rivals; while pinching, males are also trying to lift rivals off the ground and shove them away (Dennenmoser & Christy, 2013). In other species of crustaceans, such as crayfish, claws are used not only to pinch but also to push rivals (Graham *et al*., 2020). Perhaps because of this variation, weapon function has seldom been comparatively investigated across different taxa. In the few groups this has been done (e.g., bovids, Caro *et al*., 2003) such variation has been eliminated by linking fighting style of a species to a single behaviour (and discarding displays).

However, since the behaviour adopted may change, ascribing fighting style to a single behaviour may underestimate the importance of the weapon for contests success and restrict our comprehension about the selective forces that act on weapon evolution. For this reason, here we reviewed the behaviours adopted during the fight for the species selected in our meta-analysis, which include both vertebrates and invertebrates. We also identified if there are groups of species that show similar weapon functions based on more complete descriptions of fighting behaviour.

### (1) Contest descriptions

For each species selected in our meta-analysis, we searched for information on weapon function. When the selected article did not provide descriptions of fighting behaviour, we searched for additional articles that contained detailed descriptions of the fighting behaviour (Supplementary File S2). Based on the exact descriptions provided in the literature, we were able to identify general categories of how contests begin and how the weapon is used during physical contact (i.e., fighting style).

#### (a) How do contests begin?

We classified species in three groups according to how contests begin. First, we found species in which individuals perform behaviours that allow them to gather information about the rival’s body size before engaging in contests. For instance, in the cricket *Melanotes ornata,* males lash their antennae toward the rival’s body before deciding whether to use their legs to kick the rival (Lobregat *et al*., 2019). In the lizard *Carinascincus microlepidotus*, on the other hand, males turned sideways towards the rival and performed dorso-lateral displays of the body before escalating to physical contact (Olsson & Shine, 2000). We named this category ‘body size estimation’ because individuals had a chance to assess their rivals body given their displays and antennae touch, but they did not necessarily have threat displays exclusively involving the weapon (following Számadó, 2008). Second, we found species in which individuals perform behaviours that allow them to gather more precise information about their rivals’ weapons before engaging in physical contact. For example, in the crayfish *Cherax dispar*, individuals display their enlarged front claws (which is used as a weapon) to one another before making physical contact, which allow a visual estimation of weapon size (Wilson *et al*., 2007). Grasshoppers also used similar behaviours in which they flare their mandibles to the rivals before engaging in physical contact (Umbers *et al*., 2012). We named this category ‘weapon display’ because rivals had the opportunity to assess the size of the rival’s weapon (or be threatened by the rival’s weapon, see Számadó, 2008). Third, there were species that did not perform explicit behaviours that would allow rivals to gather information about body or weapon size before engaging in physical contact. An example is the *Sagra femorata* beetle, in which rivals do not use any type of display before beginning a physical struggle (Katsuki *et al*., 2014; O’Brien & Boisseau, 2018). We named this category ‘no display’ because there was no evidence that rivals gather information or threaten each other before contests ensue. Although we cannot exclude the possibility that rivals gather information during the physical contact phase, we assure that the decision to begin a contest in the group of ‘no display’ is little or not affected by mutual evaluations performed before rivals engage in a physical struggle. The full descriptions of contest behaviours for all species can be checked in Supplementary File S2.

#### (b) How are weapons used during physical contact?

We identified six categories which relate to fighting style: (i) to lift; (ii) to push; (iii) to pull; (iv) to squeeze; (v) to deliver forceful impact blows; (vi) to pierce. Lifting weapons were mainly used to lift the rival from the substrate to either disbalance or topple the rival. For instance, the stag beetle *Cyclommatus metallifer* uses enlarged mandibles to hold and lift the rival off the tree trunk in which they frequently fight on (Goyens *et al*., 2015). Pushing weapons were used to push the rival away from the bearer. For instance, dung beetles that fight inside tunnels use their horns to push rivals off the entrance of the tunnels (McCullough & Simmons, 2016). Pulling weapons were used to pull the rival near the bearer, frequently dislodging it. For instance, *Aegla longirostri* freshwater anomurans use their claws to pinch and pull the rivals, dislodging them from the substrate (Ayres-Peres, Araujo, & Santos, 2011). This behaviour is rarely performed without squeezing. Squeezing weapons were mainly used to provide forceful grasp on a rival. Crustaceans and lizards are the frequent examples of this category, using their claws and jaws to squeeze rival’s body parts (Husak, Lappin, & Van Den Bussche, 2009; Dennenmoser & Christy, 2013). Impact weapons were used to deliver rapid or explosive strikes to the rival. A noteworthy example is the raptorial appendages of mantis shrimps, which are used to strike the rival’s telson with a movement so fast that it can crack the abdomen’s cuticle (Green & Patek, 2015 but see Taylor & Patek, 2010). Lastly, piercing weapons were used mainly to pierce the rival’s skin or cuticle, typically with sharp, pointy structures. One example is the bird *Ficedula hypoleuca* which uses its beak to pierce the rival’s skin during physical contact (Dale & Slagsvold, 1995).

It is important to note that, despite these six different functions, most species used their weapons for two or, more rarely, three functions during the fight. For example, in the cricket *Loxoblemmus doenitzi,* males have a flat head with horns on the edges that are used during fights to push one another. However, males may also use the horns to rapidly beat the rival’s horns or body (Kim, Jang, & Choe, 2011). Therefore, for species in which more than one weapon usage was described, we created categories with all the functions associated with that weapon. In our example, we consider that males of *L. doenitzi* use their weapons for both pushing and impacting their rivals.

### (2) Results

Based on the combination between how contests begin and how weapons are used during contests we identified 16 categories of weapon function distributed among the 52 species (Table 3). According to the descriptions we gathered, contests frequently began by individuals displaying their weapons to rivals (n = 32 species out of 52, 61.53%); less frequently by displaying their body size (n = 17 of 52 species, 32.69%); and rarely by not making any display (n = 3 of 52 species, 5.76%). Regarding function during fights, most weapons are used for more than one function (n = 36 of 52, 69.23%), while few are used for a single function (n = 16 of 52, 30.76%). Regardless of whether we count multifunctional weapons, or weapons with a single function, squeezing is the most common function (35), followed by pushing (23), lifting (15), impacting (15), pulling (3), and piercing (2). To see how each species was categorized, see Fig. 1; for complete descriptions, see Table 3.

**Table 3.** Functions (lines) and displays (columns) of fighting styles found across the 52 species of animals used in our study. All possible combinations of the five functions were considered, even though some of them were absent for our sample. To see how each species was categorized, please see Fig. 1 and Supplementary File S2 for the descriptions.

| FUNCTION | WEAPON DISPLAY | BODY SIZE ESTIMATION | NO DISPLAY | TOTAL |
| --- | --- | --- | --- | --- |
| Squeezing | 12 | 1 | - | 13 |
| Impact | 1 | 1 | - | 2 |
| Pushing | 1 | - | - | 1 |
| Lift + Squeezing | 1 | 1 | 1 | 3 |
| Lift + Impact | 3 | - | 1 | 4 |
| Lift + Pushing | 2 | 1 | 1 | 4 |
| Squeezing + Impact | 3 | - | - | 3 |
| Squeezing + Piercing | - | 1 | - | 1 |
| Squeezing + Pushing | 1 | 5 | 1 | 7 |
| Squeeze + Pull | 1 | 2 | - | 3 |
| Impact + Pushing | - | 3 | - | 3 |
| Piercing + Pushing | 1 | - | - | 1 |
| Push + Impact + Lift | 2 | - | - | 2 |
| Push + Impact + Squeeze | 1 | 1 | - | 2 |
| Push + Lift + Squeeze | 1 | 1 | - | 2 |
| Push + Squeeze + Pull + Lift | 1 | - | - | 1 |
| **TOTAL** | **31** | **18** | **3** | **52** |

### (3) Discussion

According to contest theory, displays should be favoured in animals that frequently engage in contests as a mechanism to decrease the costs of aggression (Emlen, 2008). By displaying (the weapon or the body), an individual might induce the rival to give up on the contest even before they started - thus saving energy. That is why displays should be favoured across all animals: they decrease the likelihood of injuries and increase the amount of energy saved (Hardy & Briffa, 2013). Our result is the first to corroborate the theoretical prediction for a directional selection for displays using a diverse group of animals: we showed that displays are commonplace among fighting animals. That seems especially true for weapon displays, which is well distributed among all functions (Table 3) and hence does not seem subject to oversampling of any given taxa. Further, displaying the weapon is more common than body displays. Therefore, we have shown that selection seems to be acting to decrease the costs of aggression across animals by favouring contests that begin with displays, rather than instantaneous aggression (Emlen, 2008).

In the three cases that we found no displays, three opposite explanations arise. First, animals might escalate all contests if resource value is extremely high. In the case of the male hermit crabs *Diogenes nitidimanus*, for instance, males guard the females by grasping the outer rim of her shell and carrying her around (Yoshino, Koga, & Oki, 2011). If that male loses the possession of the female in a contest, it is likely it will not mate until the next reproductive cycle. Access to mature females is thought to be difficult in *D. nitidimanus* because females are only receptive during very short time windows (only after molting, Asakura, 1987). Thus, males holding females might opt to go all-in in a fight to keep the female, similar to predictions from the ‘Desperado’ effect (Grafen, 1987). Second, if the costs of fighting are very low, there might be no selection to avoid fighting and decrease the costs. However, the three species in our sample (*Trypoxylus dichotomus*, *Sagra femorata*, *D. nitidimanus*) have enlarged weapons (i.e., disproportional size in relation to body size, Yoshino *et al*., 2011; Johns *et al*., 2014; O’Brien, Katsuki, & Emlen, 2017). Thus, it is unlikely that these species are evolving weapons because contests have very low costs. The third explanation resides on a possible lack of details in behavioural descriptions of fighting behaviour. Those might be unconscious biases, such as focusing on what animals are doing during the fight and not when the fight is starting, or subtle signalling behaviours that are outside the observer’s sensory cognition. For instance, rhinoceros beetles can use acoustic signals during courtship that were unknown until recently (Hunt *et al*., 2020). Thus, it is possible that these species indeed use displays, but we were not able to assess them yet.

Squeezing was the most common behaviour, followed by pushing. While squeezing can be considered a bias because of the oversampling of crustaceans and lizards (both with squeezing weapons), pushing is also common behaviour during fights, although in many instances it is associated with another behaviour. As shown in Table 3, pushing was found associated with another behaviour in 31 (of 34 records), being used as the only fighting behaviour only in three species. Thus, pushing is one of the main reasons most weapons are multifunctional. It is interesting to note that, in most species, pushing is associated with squeezing – which is a similar pattern to the vectorially opposite behaviour, pulling. Individuals are unable to pull a rival without holding the rival. Pushing, however, can be done simply by contacting the individual (e.g., interlocking antlers used to push). Thus, despite these differences, it seems that squeezing might increase the chance that an individual also tries to manipulate the rival by pulling or pushing.

Piercing, on the other hand, is the rarest behaviour in our sample. The only species that displayed that behaviour in our sample was *Ficedula hypoleuca*, a bird that pecks its rivals during fights (Dale & Slagsvold, 1995), and *Tetranychus urticae*, a mite species in which individuals use their stylets to pierce rivals (Potter, Wrensch, & Johnston, 1976). Other species, such as hummingbirds (Rico-Guevara & Araya-Salas, 2015) and coreid bugs (Emberts *et al*., 2021), can use piercing as their fighting style, but even among them few groups use piercing. Piercing is indeed expected to be rare because it might be injurious. Over evolutionary time, theory predicts that species would tend to evolve displays to decrease the likelihood of engaging in injurious contests, unless resource value is extremely high (Hardy & Briffa, 2013). By evolving displays, weapons would then tend to increase in size and complexity, which could decrease piercing performance and change the function of the weapon altogether (Emlen, 2008; Anderson, 2018). A similar route is believed to have occurred in cervids, where short, pointy antlers started to increase in size and complexity as species evolved (Barrette, 1977; Caro *et al*., 2003; Davis, Brakora, & Lee, 2011). Our results show that using displays before fights is indeed a common strategy among animals (Table 3), but we still need to test whether these displays decrease the injury capacity of weapons over evolutionary time.

Another important pattern is that the fighting style of most species (65%) are comprised of multiple behaviours. Since fights tend to involve ritualized behaviours (Hardy & Briffa, 2013), it seems improbable that the description of a weapon being used in more than one behaviour occurred by chance. Therefore, it seems that there is a higher tendency for weapons to be used for more behaviours. Perhaps more complex fighting styles increase the winning chances because it gives more options for individuals to inflict costs on their rivals. But at the same time, depending on the combinations of behaviour, this may also generate opposing selective forces on weapon morphology. For example, weapons used for lifting and pushing will probably favour a single morphological type that provides an efficient way to fit the weapon on the rival and a strong body to work as a lever for the weapon to work properly. Weapons used for impacting and pushing, on the other hand, should favour a strong structure that is capable of delivering high forces, but a different morphology for pushing the rival. Perhaps, the occurrence of opposite selective forces acting on weapons explains why some functional combinations are not described (Corn *et al*., 2021). Further, if some behaviours tend to involve multiple non-weapon parts (e.g., lifting uses the legs and body), while others are essentially a weapon movement (e.g., squeezing), perhaps these behaviours should not be considered equally when weapon evolution is concerned. On the one hand, the distinct selective forces on some functions might promote weapon diversification (Wainwright *et al*., 2005; Polly, 2020). On the other hand, the weapon is primarily used solely in some behaviours. The full breadth of these possibilities remains to be investigated.

### V. DOES DISPLAY AND FIGHTING STYLE INFLUENCE WHICH WEAPON COMPONENTS ARE IMPORTANT FOR CONTEST RESOLUTION?

As shown in the previous section, species differ in how they begin contests and in how they use their weapons during fights. Given this variation, it is possible that displays and fighting style affect which traits are more important to increase contest success. In particular, it may be that, in species that display their weapons before engaging in physical contact, greater weapon size increases the chance of contest success because weapon size would deter most rivals from fighting. For species that instead assess body size, not weapon size, differences in weapon size might have a more important role during fight than before physical contact ensues. Therefore, in species that use body size displays, the difference between winners and losers in weapon size should be lower than in species that use weapon displays. Finally, in species that do not use any type of display before fighting, weapons may still be important in determining victory. The absence of a display prior to physical contact may relax the selective pressure on weapon size, but may increase the selection acting on performance depending on fighting style.

Regarding fighting style, it is possible that differences in how a weapon is used affects the relative importance of weapon size on contest resolution. For species that use weapons to lift or push rivals, reach should be important to decide who wins the contest. Basically, larger weapons allow the individual to touch its rival before being touched. That allows the individual to attack without being exposed to a riposte. Thus, we can expect a large selective pressure on the size of the weapon for these two types of fighting (up to a certain mechanical limit, see McCullough, 2014). Squeezing, impact, or piercing, on the other hand, do not necessarily rely on size. Although weapon size may still be important (to attack first and due to allometric effects, Pélabon *et al*., 2014), a larger weapon may not equate to a weapon that performs better. Crayfish, for instance, bear large claws that can be relatively weak for their overall size (Robinson & Gifford, 2019). Because weapon size may not determine its performance, it is possible that an individual with a smaller but stronger weapon can cause more injuries than an individual with a larger but weaker weapon. Therefore, we expect the mean difference in weapon size between winners and losers for weapons used to squeeze, impact, or pierce to be low.

### (1) Methods

To analyse the differences in the type of display and fighting style, we performed two multilevel meta-analytical models using only the effect sizes for linear measurements. As can be seen in section III, the data on performance measures contain only Crustacea and Squamata (and one cricket), which biases the types of display and fighting style we observe. Thus, we used the type of display evaluated in the previous session as our moderator, the Hedges’ *g* effect size as our response variable, and the inverse of the variance of Hedges’ *g* as a weight. The rest of the model, such as its random effects, and how we assessed heterogeneity, are equal to the procedures described in session 3.1.

For the fighting style analysis, we used the descriptions on the previous section to categorize fighting style in three groups: (i) Size-emphasis; (ii) Performance-emphasis; (iii) Intermediate. The size-emphasis group consisted of weapons used to pull, push and lift rivals, including weapons with these two functions. The performance-emphasis group consisted of weapons used to squeeze, impact, pierce, or pull rivals. Again, if a weapon had two of these three behaviours simultaneously, we categorized it as ‘performance-emphasis’. Any weapon used for two or more behaviours that belonged to two different groups (i.e., ‘size’ and ‘performance-emphasis’), was categorized as ‘intermediate’ (all species that used the weapons for three behaviours were included in this last category). For this multilevel meta-analytical model, we used the function group as our moderator. We used the same random effects and heterogeneity assessments described in session 3.1.

### (2) Results

#### (a) How do fights begin?

Winners had larger weapons than losers regardless of the type of display on average (QM_3_ = 28.48, p < 0.0001, Fig. 3), but the confidence interval overlapped zero when males did not use displays (Fig. 3). Further, the types of displays did not differ among themselves (QM_2_ = 3.18, p = 0.2). The model showed low heterogeneity. Study ID and pairing method were responsible for most of that heterogeneity, while phylogeny and genus had negligible effects on variance (Table 4). We found evidence for publication bias on the effect sizes (Egger’s test; intercept: -0.368, 95% CI: -0.656, -0.081, t = -2.553, p = 0.012).

**Fig. 3.**
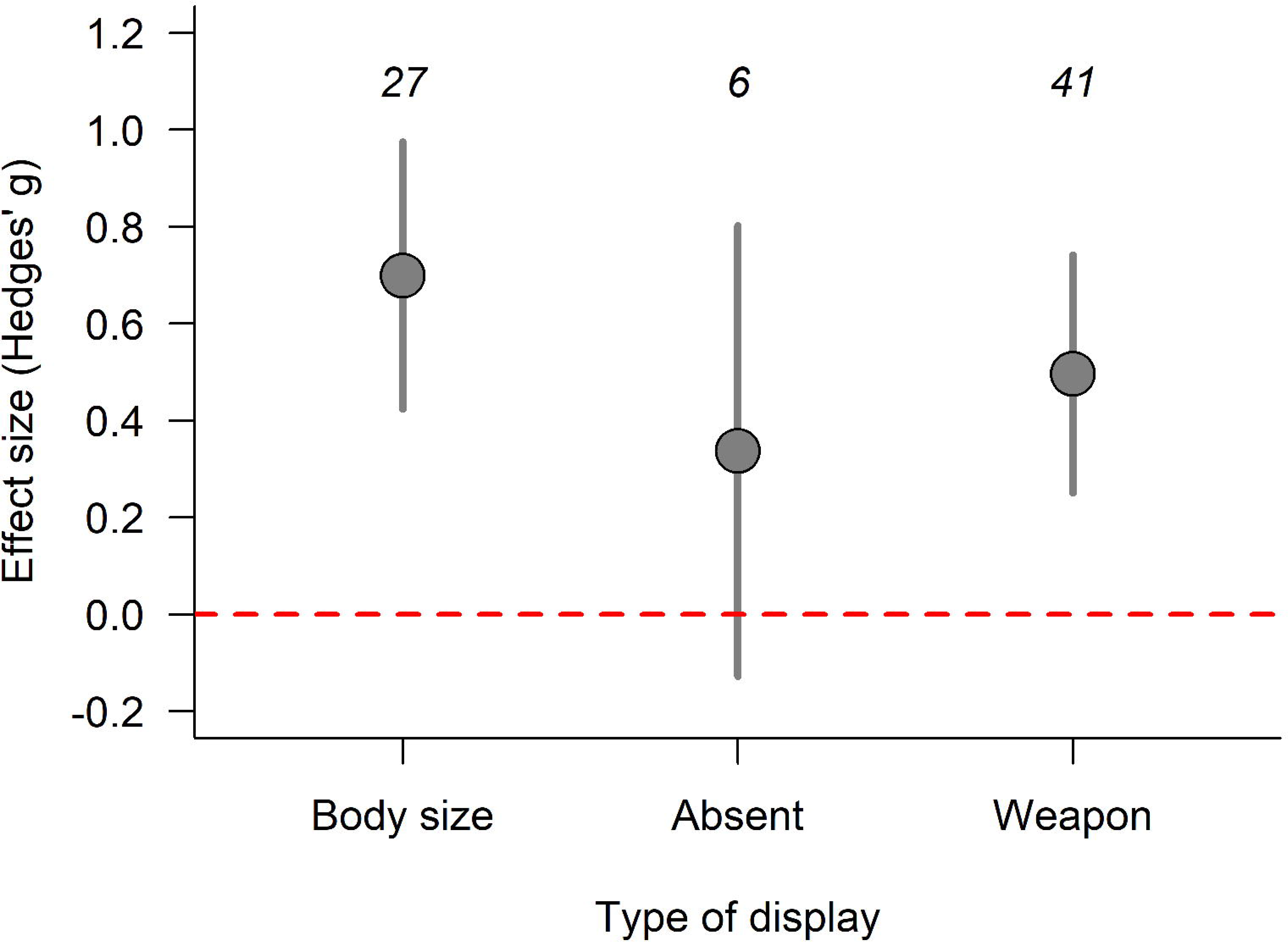
Winners had larger weapons than losers regardless of the type of display, but these differences were similar among the types of display. The effect size, Hedges’ *g*, represents the mean standardized difference in linear measures between winners and losers. Positive values denote that winners had larger weapons than losers, while negative values represent the opposite. Dots represent the estimated values from a multilevel meta-analytic model considering Hedges’ *g* as the response variable and the type of display as a moderator variable. The error bars represent the 95% confidence interval of the estimate. The numbers above the error bars represent the number of effect size in each component.

**Table 4.** Heterogeneity of the meta-analytical model of differences in how contests begin (i.e., if displays are used, and which type of displays).

| Random effect | I<sup>2</sup> (95% CI) |
| --- | --- |
| Study ID | 9.75 (7.83, 11.67) |
| Pairing method | 17.93 (16.01, 19.85) |
| Phylogeny | 5.84 (3.92, 7.76) |
| Genus | < 0.0001 (0.00, 1.92) |
| Total | 33.54 (31.62, 35.46) |
| *H*<sup>2</sup> | 0.17 (0.00, 2.09) |

#### (b) How do weapons differ among fighting style?

Winners were larger than losers in all categories, even though some of them had confidence intervals that slightly overlapped with zero (QM_3_ = 107.72, p < 0.0001, Fig. 4). We also found differences in the asymmetry between winners and losers among the categories. Weapons were more important for contest success in size-emphasis and intermediate fighting style when compared to performance-emphasis (QM_1_ = 18.84, p < 0.001, QM_1_ = 8.42, p = 0.003, respectively). The model showed low heterogeneity. Study ID and pairing method were responsible for most of that heterogeneity, while phylogeny and genus had negligible effects on variance (Table 5). We found evidence for publication bias on the effect sizes (Egger’s test; intercept: -0.368, 95% CI: -0.656, -0.081, t = -2.553, p = 0.012).

**Fig. 4.**
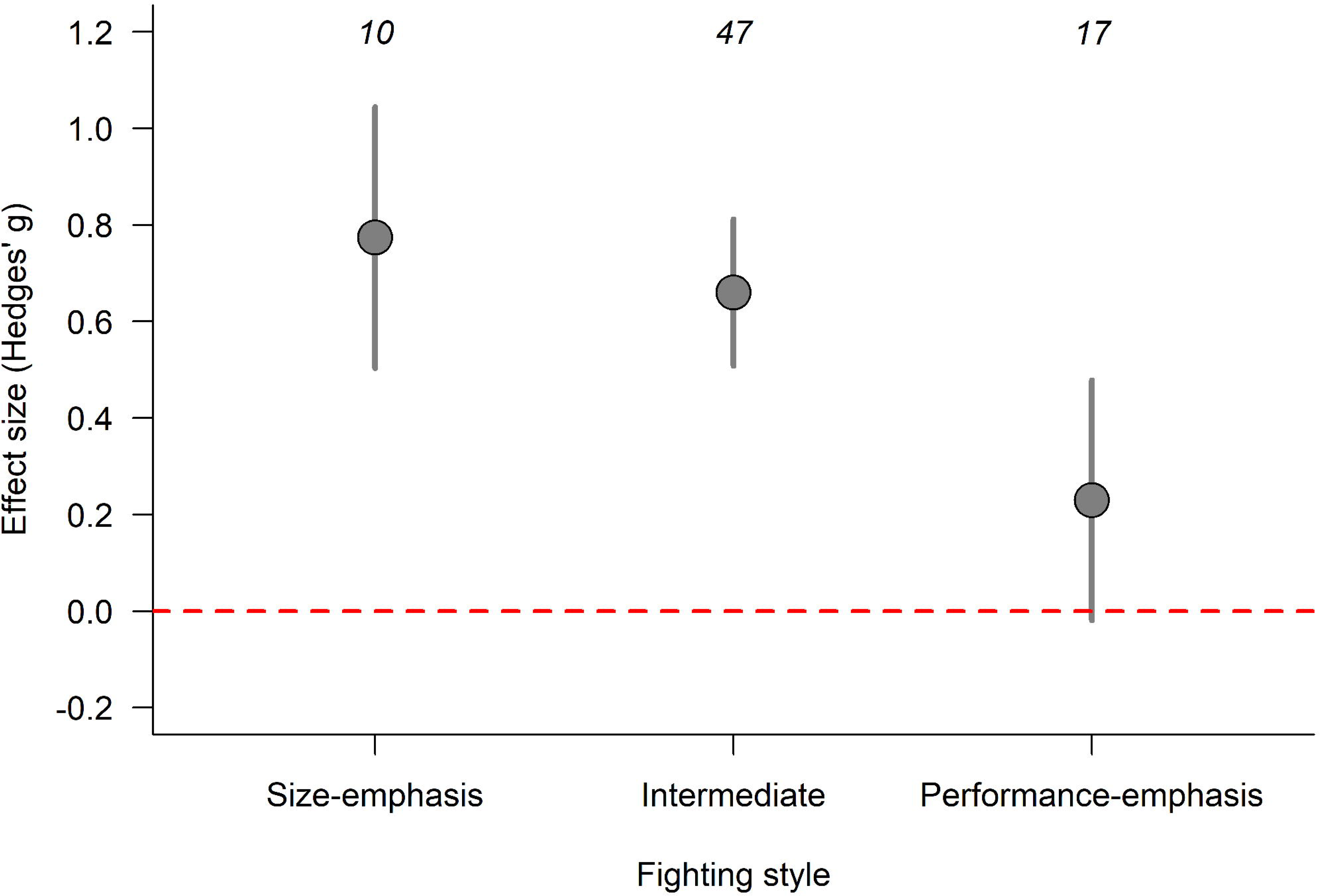
Winners had weapons that were, on average, larger than losers; and asymmetries between winners and losers in performance-emphasis were lower than the other two fighting styles. The effect size, Hedges’ *g*, represents the mean standardized difference in the linear measures of weapons between winners and losers. Positive values denote that winners had larger weapons than losers, while negative values represent the opposite. Dots represent the estimated values from a multilevel meta-analytic model considering Hedges’ *g* as the response variable and the fighting style as a moderator variable. The error bars represent the 95% confidence interval of the estimate. The numbers above the error bars represent the number of effect size in each component.

**Table 5.** Heterogeneity and variance of the meta-analytical model of differences in how weapons are used during contests.

| Random effect | I<sup>2</sup> (95% CI) |
| --- | --- |
| Study ID | 6.91 (4.99, 8.83) |
| Pairing method | 18.46 (16.54, 20.38) |
| Phylogeny | < 0.0001 (0.00, 1.92) |
| Genus | < 0.0001 (0.00, 1.92) |
| Total | 25.38 (23.46, 27.30) |
| *H*<sup>2</sup> | < 0.0001 (0.00, 1.92) |

### (3) Discussion

Contrary to our expectations, there was no difference in weapon size between winners and losers among different types of display. Therefore, the existence of a weapon or body display before the physical contact phase does not seem to impose additional selective pressures on weapon size. This is interesting because visual displays are thought to be one important driver for increments in weapon size (O’Brien *et al*., 2017; Eberhard *et al*., 2018), and most of the species in our sample perform visual displays (Fig. 3).

On the other hand, the fighting style seems to impose different pressures on weapon evolution: the mean difference between winners and losers was higher for the “size-emphasis” and “intermediate” groups when compared with the “performance-emphasis” group. Therefore, it seems that, even if a weapon is also used for squeezing, piercing or impacting, the presence of an additional behaviour related to lifting or pushing will favour increases in weapon size. Perhaps lifting and pushing are important in multifunctional weapons because they allow an individual to reach the rival before being reached or to physically expel a rival from the defended resource. For instance, *Melanotes ornata* crickets, and fiddler crabs can use their weapons to throw their rivals away from the contested resource (Supplementary File S2).

It is important to highlight, however, that weapons have many different aspects that may affect how they impose costs on rivals during the contest (Palaoro & Briffa, 2017). Therefore, to reach a more comprehensive picture of how different weapon traits change according to fighting style it is necessary to gather information on many different weapon aspects - and many different weapon types. What is clear from our results is that species show marked differences in fighting style and that such differences have the potential to affect (at least for size) the way weapons evolve.

## VI. CONCLUDING REMARKS

1. Our data on 52 species of both vertebrates and invertebrates shows biases on how animal weapons are measured. Linear components, such as lengths and widths, are responsible for 69.15% of the information on the literature; performance components, such as bite force, for 21.49%; and any other measures such as weapon asymmetry, mass, or volume are rare.
2. We also found significant bias on the taxa in which performance components are measured. Lizards and crustaceans (and one cricket species) are the only taxa in which we found measures of weapon performance (e.g., bite force) associated with winning or losing fights. Thus, there is a significant gap on how we perceive the importance of weapon performance to solve contests. To have a better idea of how performance is important, we need more information on the performance of weapons such as horns, antlers, and spines.
3. Linear components of weapons were more associated with victory than performance components. This can be a corollary of the bias mentioned above, but our sensitivity analysis suggests that the effect is robust. One possible explanation is that size can allow individuals to handle rivals from a safe distance, regardless of the weapon type. Handling rivals from a safe distance can give the individual more time to get a tighter grip and increase their chance of winning the contest.
4. The majority of species tend to begin contests by displaying their weapons to the rival (59.61%); a smaller portion either displays their body through movements or touch the rival with antennae or any other mechanosensory morphology (34.61%). A minority begins contests with no display at all (5.76%). Hence, as theory suggests, most species do begin contests by using displays.
5. Few weapons were used for a single behaviour during contests (30.8%), most of them were used for two or more behaviours (69.2%). From those multiple behaviours, pushing rivals with the weapon is the most frequent co-occurring behaviour, suggesting that selection may not be working solely on a weapon’s sole behaviour, but rather on the fighting style. Further, piercing is the rarest behaviour in our sample with only two species described with those weapons.
6. We found no strong effect of type of display on the importance of weapon size on contest resolution. The only subtle difference we found was that, when displays were absent, weapon size did not differ between winners and losers. Given that most species in our sample perform some type of display, it suggests that displays may signal weapon size. However, we need more data from species that do not perform any type of display to understand the effect properly.
7. Fighting style influences the importance of weapon size on contests. Weapons used to squeeze, impact or pierce had a lower difference between winners and losers when compared to weapon used for push, pull, or lift or weapons with multiple behaviours. Once again, it seems that being able to reach the rival first is important for contest success.

## Acknowledgments

We thank Dr. Patrick Green for the suggestions on an earlier version of the manuscript. We thank all the authors who shared the raw data of their paper with us: Chatchawan Chaisuekul, Christine W. Miller, Devin O’Brien, Kate Umbers, Leilani Walker, Marcelo M. Dalosto, Murray Fea, Sarah E. Reece, Zach Stahlschmidt, Zachary Emberts, and Zackary Graham. We also thank Eduardo S.A. Santos for his help with the heterogeneity and bias analyses. A.V.P. thanks FAPESP for the post-doctoral grant (process no: 2016/22679-3). P.E.C.P. thanks Conselho Nacional de Desenvolvimento Científico e Tecnológico (CNPq produtividade em pesquisa 311212/2018-2) and Pesquisa e Desenvolvimento of Agência Nacional de Energia Elétrica and Companhia Energética de Minas Gerais (P&D ANEEL/CEMIG, PROECOS project GT-599).

## Authors’ contribution

A.V.P and P.E.C.P. designed the study, A.V.P. collected the data and ran the analysis, A.V.P. and P.E.C.P. wrote the manuscript.

## Data availability

All data and codes will be shared on github (https://github.com/alexandrepalaoro) upon acceptance.

