## Supplementary material for "The importance of animal weapons and fighting style in animal contests": Table S1

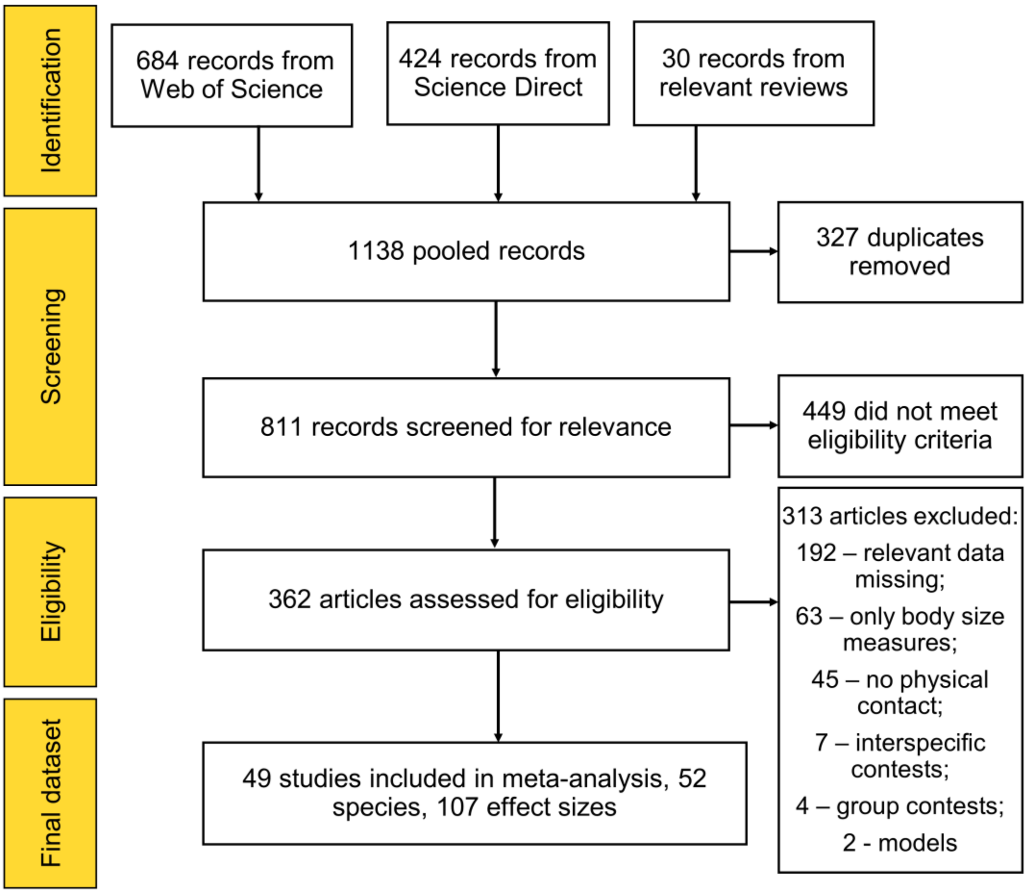


**Fig S1**. Systematic review flow diagram (PRISMA diagram, Liberati et al. 2009) of how we identified, screened, and checked eligibility of all the papers we found during our literature review.

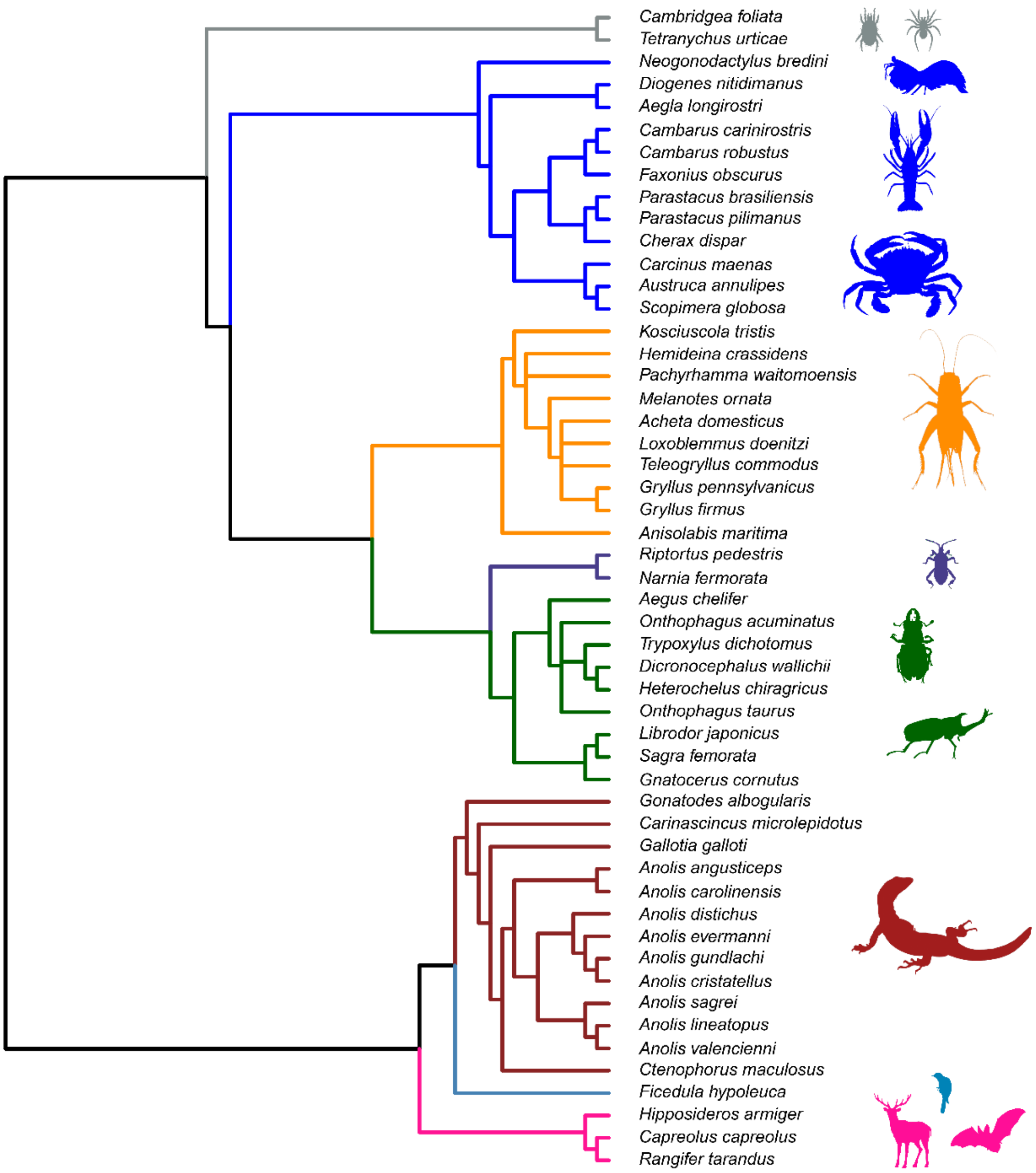


**Fig S2.** Phylogenetic tree used to estimate phylogenetic relatedness in our meta-analytic models. Since we did not have species of the same genus, we used genus tree tips. Colours indicate groups within the same order. Tree built using iTOL (https://itol.embl.de/).

| Species | Pairing | Measure | Display | Fighting style | Study | Hedges *g* [95% CI] |
|---|---|---|---|---|---|---|
| *Teleogryllus commodus* | Paired | Performance | Weapon | Impact & Lift | 45 | 0.43 [−0.02, 0.89] |
| *Pachyrhamma waitomoensis* | Random | Linear | Body | Push & Impact | 24 | 1.00 [ 0.57, 1.44] |
| *Melanotes ornata* | Random | Linear | Body | Push & Impact | 16 | 0.64 [ 0.18, 1.09] |
| *Loxoblemmus doenitzi* | Random | Linear | Body | Push & Impact | 3 | 0.36 [−0.32, 1.04] |
| *Kosciuscola tristis* | Random | Linear | Weapon | Squeeze | 30 | 0.27 [−0.22, 0.75] |
| *Hemideina crassidens* | Random | Linear | Weapon | Pull & Squeeze | 9 | 1.03 [ 0.44, 1.62] |
| *Gryllus pennsylvanicus* | Paired | Linear | Weapon | Push & Squeeze | 40 | 0.81 [ 0.13, 1.49] |
| *Gryllus pennsylvanicus* | Paired | Linear | Weapon | Push & Squeeze | 40 | 0.67 [ 0.15, 1.19] |
| *Gryllus pennsylvanicus* | Paired | Linear | Weapon | Push & Squeeze | 39 | 0.24 [−0.21, 0.68] |
| *Gryllus pennsylvanicus* | Paired | Linear | Weapon | Push & Squeeze | 39 | 0.22 [−0.22, 0.66] |
| *Gryllus pennsylvanicus* | Paired | Linear | Weapon | Push & Squeeze | 39 | 0.25 [−0.20, 0.70] |
| *Gryllus pennsylvanicus* | Paired | Asymmetry | Weapon | Push & Squeeze | 39 | 0.00 [−0.44, 0.44] |
| *Gryllus firmus* | Random | Linear | Weapon | Lift | 28 | 0.53 [ 0.29, 0.77] |
| *Acheta domesticus* | Random | Linear | Weapon | Impact & Lift | 1 | −0.05 [−0.53, 0.43] |
| *Acheta domesticus* | Random | Linear | Weapon | Impact & Lift | 1 | 0.29 [−0.19, 0.78] |
| *Rangifer tarandus* | Random | Linear | Weapon | Push, Impact & Lift | 19 | 3.29 [ 1.25, 5.33] |
| *Rangifer tarandus* | Random | Linear | Weapon | Push, Impact & Lift | 19 | 1.38 [ 0.48, 2.28] |
| *Hipposideros armiger* | Random | Linear | Body | Impact | 33 | 0.09 [−0.26, 0.44] |
| *Capreolus capreolus* | Random | Index | Weapon | Push, Impact & Lift | 4 | 2.34 [ 1.38, 3.30] |
| *Gonatodes albogularis* | Random | Linear | Weapon | Squeeze | 10 | 0.36 [−0.24, 0.96] |
| *Gonatodes albogularis* | Random | Linear | Weapon | Squeeze | 10 | 0.08 [−0.51, 0.68] |
| *Gallotia galloti* | Random | Linear | Weapon | Squeeze | 11 | 0.28 [−0.35, 0.90] |
| *Gallotia galloti* | Paired | Performance | Weapon | Squeeze | 49 | 0.36 [ 0.09, 0.63] |
| *Gallotia galloti* | Paired | Linear | Weapon | Squeeze | 47 | 0.69 [ 0.11, 1.28] |
| *Gallotia galloti* | Paired | Linear | Weapon | Squeeze | 47 | 0.02 [−0.49, 0.54] |
| *Ctenophorus maculosus* | Paired | Performance | Weapon | Squeeze | 42 | 0.41 [ 0.07, 0.75] |
| *Ctenophorus maculosus* | Paired | Linear | Weapon | Squeeze | 42 | 0.32 [−0.02, 0.66] |
| *Carinascincus microlepidotus* | Random | Linear | Body | Squeeze | 13 | −0.09 [−0.61, 0.43] |
| *Carinascincus microlepidotus* | Random | Linear | Body | Squeeze | 13 | 0.21 [−0.31, 0.74] |
| *Anolis valencienni* | Paired | Performance | Weapon | Squeeze | 41 | 0.19 [−0.49, 0.86] |
| *Anolis sagrei* | Paired | Performance | Weapon | Squeeze | 41 | 0.47 [−0.11, 1.05] |
| *Anolis lineatopus* | Paired | Performance | Weapon | Squeeze | 41 | 0.36 [−0.25, 0.97] |
| *Anolis gundlachi* | Paired | Performance | Weapon | Squeeze | 41 | 0.43 [−0.23, 1.10] |
| *Anolis evermanni* | Paired | Performance | Weapon | Squeeze | 41 | 0.23 [−0.34, 0.80] |
| *Anolis distichus* | Paired | Performance | Weapon | Squeeze | 41 | −0.35 [−0.95, 0.25] |
| *Anolis cristatellus* | Paired | Performance | Weapon | Squeeze | 41 | 0.24 [−0.35, 0.84] |
| *Anolis carolinensis* | Paired | Performance | Weapon | Squeeze | 41 | 0.51 [−0.14, 1.16] |
| *Anolis angusticeps* | Paired | Performance | Weapon | Squeeze | 41 | 0.44 [−0.19, 1.07] |
| *Neogonodactylus bredini* | Paired | Performance | Weapon | Impact | 44 | −0.04 [−0.51, 0.44] |
| *Riptortus pedestris* | Random | Linear | Body | Push, Squeeze & Impact | 17 | 1.75 [ 1.18, 2.32] |
| *Narnia femorata* | Random | Linear | Weapon | Impact & Squeeze | 27 | 0.57 [−0.85, 1.99] |
| *Narnia femorata* | Random | Linear | Weapon | Impact & Squeeze | 27 | 0.69 [−0.75, 2.13] |
| *Narnia femorata* | Random | Linear | Weapon | Impact & Squeeze | 27 | 0.03 [−0.85, 0.92] |
| *Narnia femorata* | Random | Linear | Weapon | Impact & Squeeze | 27 | 0.10 [−0.78, 0.99] |
| *Narnia femorata* | Random | Linear | Weapon | Impact & Squeeze | 27 | 0.42 [−0.47, 1.32] |
| *Narnia femorata* | Random | Linear | Weapon | Impact & Squeeze | 27 | 0.43 [−0.46, 1.33] |
| *Narnia femorata* | Random | Area | Weapon | Impact & Squeeze | 27 | 0.78 [−0.67, 2.23] |
| *Narnia femorata* | Random | Area | Weapon | Impact & Squeeze | 27 | 0.07 [−0.81, 0.96] |
| *Narnia femorata* | Random | Area | Weapon | Impact & Squeeze | 27 | 0.48 [−0.42, 1.38] |
| *Anisolabis maritima* | Paired | Asymmetry | Weapon | Impact & Squeeze | 48 | 0.26 [−0.74, 1.26] |
| *Anisolabis maritima* | Paired | Asymmetry | Weapon | Impact & Squeeze | 48 | 1.31 [ 0.37, 2.25] |
| *Scopimera globosa* | Random | Linear | Weapon | Push | 25 | 0.85 [ 0.41, 1.29] |
| *Parastacus pilimanus* | Paired | Linear | Weapon | Push, Pull & Squeeze | 37 | −0.11 [−1.25, 1.03] |
| *Parastacus brasiliensis* | Paired | Linear | Weapon | Push, Pull & Squeeze | 37 | 0.04 [−1.02, 1.10] |
| *Faxonius obscurus* | Random | Performance | Body | Push & Squeeze | 34 | 0.76 [ 0.38, 1.14] |
| *Faxonius obscurus* | Random | Performance | Body | Push & Squeeze | 34 | −0.17 [−0.55, 0.20] |
| *Faxonius obscurus* | Random | Linear | Body | Push & Squeeze | 34 | 0.87 [ 0.49, 1.25] |
| *Faxonius obscurus* | Random | Linear | Body | Push & Squeeze | 34 | 0.86 [ 0.47, 1.25] |
| *Diogenes nitidimanus* | Random | Linear | No | Push & Squeeze | 15 | 0.46 [−0.02, 0.94] |
| *Cherax dispar* | Random | Performance | Weapon | Push & Squeeze | 14 | 0.73 [ 0.21, 1.25] |
| *Cherax dispar* | Random | Linear | Weapon | Push & Squeeze | 14 | 0.43 [−0.05, 0.91] |
| *Carcinus maenas* | Random | Linear | Weapon | Push, Squeeze & Impact | 31 | 0.45 [−0.22, 1.13] |
| *Carcinus maenas* | Random | Linear | Weapon | Push, Squeeze & Impact | 31 | 0.33 [−0.34, 1.00] |
| *Carcinus maenas* | Paired | Performance | Weapon | Push, Squeeze & Impact | 36 | 0.59 [ 0.04, 1.14] |
| *Carcinus maenas* | Paired | Mass | Weapon | Push, Squeeze & Impact | 35 | 0.41 [−0.34, 1.17] |
| *Carcinus maenas* | Paired | Mass | Weapon | Push, Squeeze & Impact | 36 | 0.34 [−0.20, 0.88] |
| *Carcinus maenas* | Paired | Linear | Weapon | Push, Squeeze & Impact | 35 | 0.61 [ 0.02, 1.20] |
| *Carcinus maenas* | Paired | Area | Weapon | Push, Squeeze & Impact | 35 | 0.50 [−0.26, 1.26] |
| *Cambarus robustus* | Random | Performance | Body | Push & Squeeze | 34 | 1.01 [ 0.54, 1.47] |
| *Cambarus robustus* | Random | Performance | Body | Push & Squeeze | 34 | −0.11 [−0.54, 0.33] |
| *Cambarus robustus* | Random | Linear | Body | Push & Squeeze | 34 | 1.81 [ 1.29, 2.34] |
| *Cambarus robustus* | Random | Linear | Body | Push & Squeeze | 34 | 1.70 [ 1.18, 2.21] |
| *Cambarus robustus* | Paired | Linear | Body | Push & Squeeze | 38 | 0.38 [−0.16, 0.92] |
| *Cambarus robustus* | Paired | Linear | Body | Push & Squeeze | 38 | 0.35 [−0.19, 0.89] |
| *Cambarus robustus* | Paired | Linear | Body | Push & Squeeze | 38 | 0.37 [−0.17, 0.91] |
| *Cambarus robustus* | Paired | Linear | Body | Push & Squeeze | 38 | 0.40 [−0.14, 0.94] |
| *Cambarus carinirostris* | Random | Performance | Body | Push & Squeeze | 34 | 0.80 [ 0.39, 1.21] |
| *Cambarus carinirostris* | Random | Performance | Body | Push & Squeeze | 34 | 0.25 [−0.15, 0.64] |
| *Cambarus carinirostris* | Random | Linear | Body | Push & Squeeze | 34 | 1.38 [ 0.95, 1.82] |
| *Cambarus carinirostris* | Random | Linear | Body | Push & Squeeze | 34 | 1.31 [ 0.88, 1.75] |
| *Austruca annulipes* | Random | Linear | Weapon | Squeeze & Lift | 8 | 0.50 [ 0.22, 0.78] |
| *Aegla longirostri* | Random | Performance | Body | Pull & Squeeze | 2 | 1.27 [ 0.47, 2.07] |
| *Aegla longirostri* | Random | Linear | Body | Pull & Squeeze | 2 | 1.22 [ 0.43, 2.01] |
| *Aegla longirostri* | Random | Linear | Body | Pull & Squeeze | 2 | 1.28 [ 0.48, 2.08] |
| *Aegla longirostri* | Paired | Performance | Body | Pull & Squeeze | 2 | −0.00 [−0.72, 0.72] |
| *Aegla longirostri* | Paired | Linear | Body | Pull & Squeeze | 2 | 0.01 [−0.71, 0.73] |
| *Aegla longirostri* | Paired | Linear | Body | Pull & Squeeze | 2 | 0.03 [−0.69, 0.75] |
| *Trypoxylus dichotomus* | Random | Linear | No | Push & Lift | 6 | 0.34 [−0.05, 0.73] |
| *Trypoxylus dichotomus* | Random | Linear | No | Push & Lift | 5 | 1.47 [ 0.76, 2.18] |
| *Trypoxylus dichotomus* | Random | Linear | No | Push & Lift | 5 | 0.22 [−0.37, 0.81] |
| *Sagra femorata* | Random | Linear | No | Squeeze & Lift | 26 | 0.45 [ 0.16, 0.73] |
| *Sagra femorata* | Random | Linear | No | Squeeze & Lift | 26 | 0.29 [ 0.00, 0.57] |
| *Onthophagus taurus* | Paired | Linear | Weapon | Push & Lift | 43 | 1.04 [ 0.49, 1.59] |
| *Onthophagus acuminatus* | Paired | Linear | Weapon | Push & Lift | 46 | 0.70 [ 0.15, 1.25] |
| *Librodor japonicus* | Random | Linear | Weapon | Push, Squeeze & Pry | 22 | 0.81 [ 0.37, 1.25] |
| *Librodor japonicus* | Random | Linear | Weapon | Push, Squeeze & Pry | 12 | 0.80 [ 0.04, 1.56] |
| *Librodor japonicus* | Random | Linear | Weapon | Push, Squeeze & Pry | 12 | 0.14 [−0.64, 0.92] |
| *Heterochelus chiragricus* | Random | Linear | Weapon | Squeeze & Lift | 29 | 0.74 [ 0.02, 1.46] |
| *Heterochelus chiragricus* | Random | Linear | Weapon | Squeeze & Lift | 29 | 0.83 [ 0.60, 1.06] |
| *Gnatocerus cornutus* | Random | Linear | Weapon | Push, Squeeze & Pry | 21 | 0.56 [ 0.08, 1.05] |
| *Dicronocephalus wallichii* | Random | Linear | Body | Push & Lift | 18 | 2.53 [ 1.24, 3.82] |
| *Aegus chelifer* | Random | Linear | Body | Push, Squeeze & Pry | 20 | 0.78 [ 0.49, 1.08] |
| *Aegus chelifer* | Random | Linear | Body | Push, Squeeze & Pry | 20 | 0.74 [ 0.44, 1.03] |
| *Aegus chelifer* | Random | Linear | Body | Push, Squeeze & Pry | 20 | 0.73 [−0.19, 1.65] |
| *Ficedula hypoleuca* | Random | Linear | Body | Squeeze & Piercing | 7 | 0.24 [−0.65, 1.12] |
| *Cambridgea foliata* | Random | Linear | Body | Push | 23 | 0.59 [ 0.24, 0.94] |
| *Tetranychus urticae* | Random | Linear | Weapon | Push & Piercing | 32 | 1.35 [ 0.22, 2.48] |

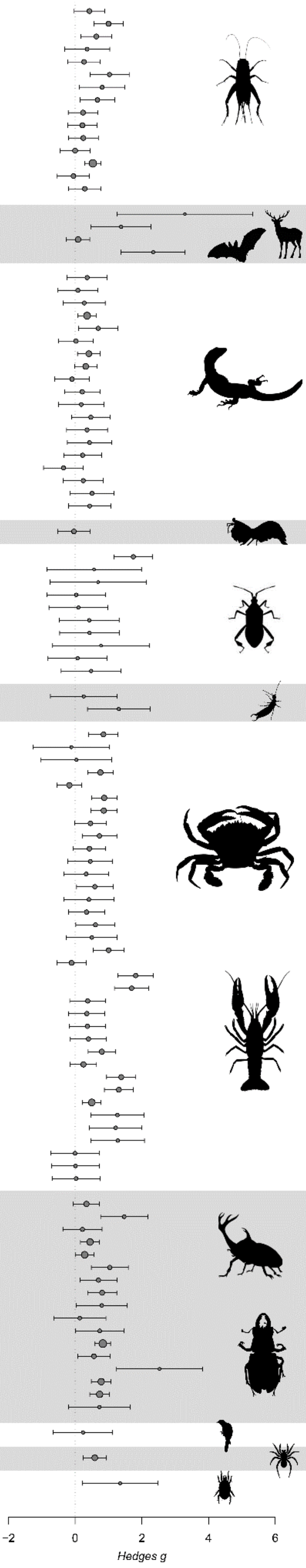

**Fig. S3.** Hedges' *g* value (circle) and corresponding confidence intervals (lines) for species, pairing method (whether contests were paired for body size or not), component measured (e.g., length, width, mass), type of display (i.e., displaying weapon, body size, or not displaying) and function of the weapon during the contests (see topic 3). Study demonstrates which samples were taken from the same study (references below). Larger grey circles denote studies with larger sample sizes. Silhouettes were taken from phylopic. Grey rows were selected randomly to facilitate visual separation of groups.

**REFERENCES FROM FIGURE 1**

1 = Briffa, 2008; 2 = Palaoro *et al.*, 2014; 3 = Kim, Jang, & Choe, 2011; 4 = Hoem *et al.*, 2007; 5 = Hongo, 2003; 6 = Karino, Niiyama, & Chiba, 2005a; 7 = Dale & Slagsvold, 1995; 8 = Jennions & Backwell, 1996; 9 = Kelly, 2006; 10 = Martínez-Cotrina, Bohórquez-Alonso, & Molina-Borja, 2014; 11 = Molina-Borja, Padron-Fumero, & Alfonso-Martin, 1998; 12 = Okada & Miyatake, 2004; 13 = Olsson & Shine, 2000; 14 = Wilson *et al.*, 2007; 15 = Yoshino, Koga, & Oki, 2011; 16 = Lobregat *et al.*, 2019; 17 = Okada *et al.*, 2011; 18 = Kojima & Lin, 2017; 19 = Barrette & Vandal, 1986; 20 = Songvorawit, Butcher, & Chaisuekul, 2018; 21 = Okada, Miyanoshita, & Miyatake, 2006; 22 = Okada & Miyatake, 2007; 23 = Walker & Holwell, 2018; 24 = Fea & Holwell, 2018; 25 = Ida & Wada, 2017; 26 = O'Brien & Boisseau, 2018; 27 = Nolen, Allen, & Miller, 2017; 28 = Nguyen & Stahlschmidt, 2019; 29 = Rink *et al.*, 2019; 30 = Umbers *et al.*, 2013; 31 = Souza *et al.*, 2019; 32 = Potter, Wrensch, & Johnston, 1976; 33 = Sun *et al.*, 2019; 34 = Graham & Angilletta, 2020; 35 = Sneddon, Huntingford, & Taylor, 1997; 36 = Sneddon *et al.*, 2000; 37 = Dalosto *et al.*, 2013; 38 = Guiasu & Dunham, 1997; 39 = Judge & Bonanno, 2008; 40 = Judge *et al.*, 2010; 41 = Lailvaux & Irschick, 2007; 42 = McLean & Stuart-Fox, 2015; 43 = Moczek & Emlen, 2000; 44 = Green & Patek, 2015; 45 = Hall *et al.*, 2010; 46 = Emlen, 1997; 47 = Bohórquez-Alonso *et al.*, 2018; 48 = Munoz & Zink, 2012; 49 = Huyghe *et al.*, 2005

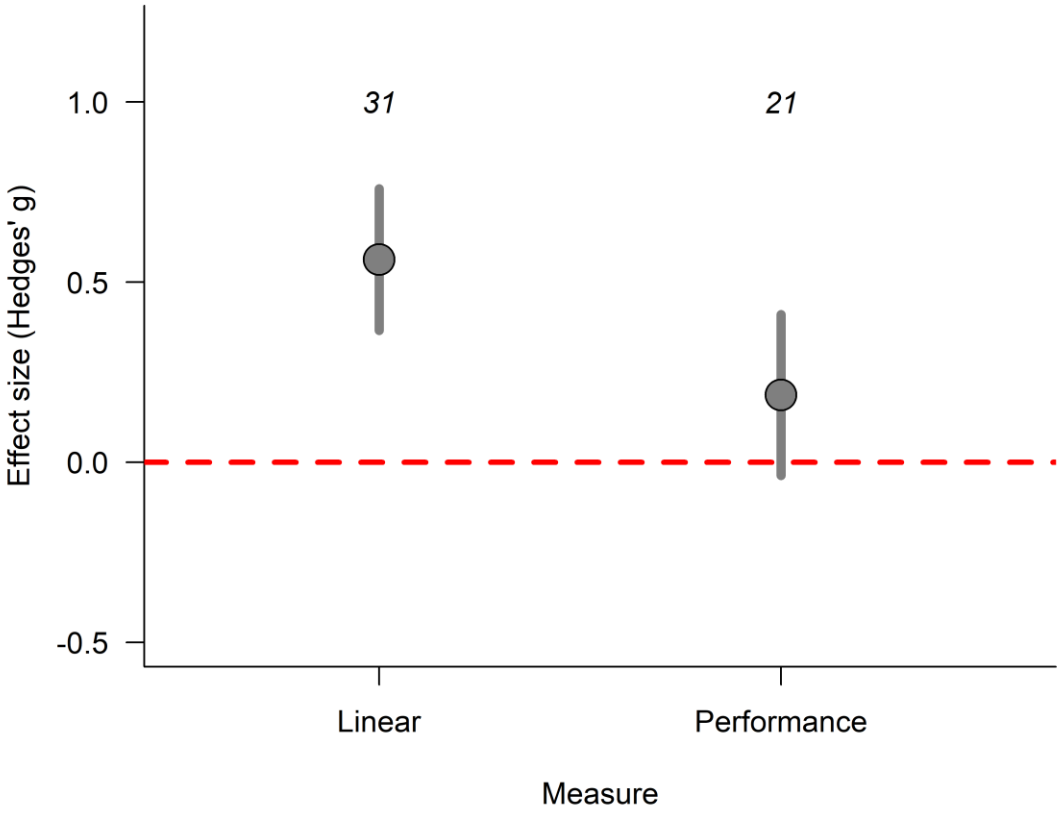


**Fig S4.** When comparing only measures taken from crustaceans and lizards, linear measures were still relatively more important to resolve contests than performance measures ($QM_1 = 18.198$, $p < 0.0001$). The effect size, Hedges' g, represents the mean standardized difference between winners and losers. Positive values denote that winners were larger than losers, while negative values represent the opposite. Dots represent the estimated values from a multilevel meta-analytic model considering Hedges' *g* as the response variable, weapon component as a moderator variable, and study ID, species ID, phylogeny, environment and pairing method as random variables. The error bars represent the 95% confidence interval of the estimate. The numbers above the error bars represent the number of effect size in each component.

**Table S1.** Species that did not have the Pearson's correlation coefficient between body size and weapon size in the same paper where the effect size was taken from . Thus, we searched for the coefficient in other papers for similar species that bore similar weapons.

| **Species** | **Reference** |
|---|---|
| *Loxoblemmus doenitzi* | Kuriwada T. (2016) Horn length is not correlated with calling efforts in the horn-headed cricket *Loxoblemmus doenitzi* (Orthoptera: Gryllidae). Entomol Sci. 19(3):228–32. |
| *Acheta domesticus* | Judge KA, Bonanno VL. Male Weaponry in a Fighting Cricket. PLoS ONE, 3(12). |
| *Gallotia galloti* | Herrel A, Spithoven L, Damme RV, Vree FD. Sexual dimorphism of head size in *Gallotia galloti*: testing the niche divergence hypothesis by functional analyses. Funct Ecol. 1999;13(3):289–97. |
| *Rangifer tarandus* | Melnycky, N. A., Weladji, R. B., Holand, Ø., & Nieminen, M. (2013). Scaling of antler size in reindeer (*Rangifer tarandus*): sexual dimorphism and variability in resource allocation. Journal of Mammalogy, 94(6), 1371-1379. |
