## Supplemental File S2 for "The importance of animal weapons and fighting style in animal contests"

SUPPLEMENTARY FILE S2

Below, you will find the descriptions of contest behaviour of the species selected for our meta-analysis. We copied the exact description of the behaviour in the original articles from which the descriptions were taken. We italicised the parts where the authors describe the display; and we bolded the descriptions of the function of the weapon. In some instances, we also added YouTube links of videos of the species fighting (which were provided by the author). If no reference is provided, the description was taken from the article used for the meta-analysis itself. Otherwise, the reference where the information is located is presented below the description.

**1.** *Loxoblemmus doenitzi*, Orthoptera:

**CLASSIFICATION**: *BODY SIZE ESTIMATION*; **IMPACT AND PUSH**

*"An agonistic encounter between males of L. doenitzi typically began with antennal contact. After antennal contact, two competing males typically engaged in either antennal fencing or production of aggressive song. Antennal fencing occurred when males rapidly antennated each others' antennae.* **This sometimes proceeded to grappling, in which two males faced head to head and pushed against each other** (Fig. 2a). **Because each male's head was flat, there was almost no gap between the heads of two males during grappling.** Two aggressive behaviors, **horn fencing and horn jab**, were often noted during the process. **Horn fencing occurred when a cricket rapidly beat the opponent's horns with his own horns** (Fig. 2b), **and horn jab occurred when cricket poked the opponent's body with his horns** (Fig. 2c). Following grappling, one of the males, the winner, chased the other, the loser.

**2.** *Pachyrhamma waitomoensis*, Orthoptera:

**CLASSIFICATION**: *BODY SIZE ESTIMATION*; **IMPACT AND PUSH**

**Male *P. waitomoensis* fought by turning backwards and engaging their hindlegs with those of their rival**, following a typical pattern of escalation from minor encounters to prolonged grappling (Fig. 4). *We were not able to identify any particular events during fights that consistently signalled when one weta would de-escalate or retreat*, and no injuries were observed during the contests. **Although weta are capable of biting each another, we did not observe any males doing so. Among all contests observed, only one resulted in a combatant being pried off the cave wall.**

(Figure taken from the paper to highlight the antennal fencing during the beginning of the contest)

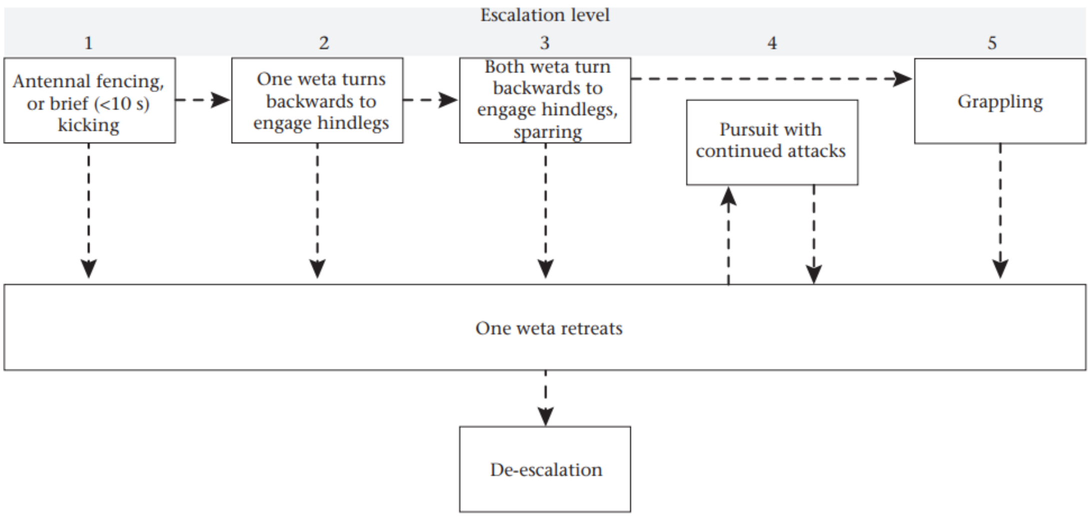


**Figure 4.** Schematic of male *P. waitomoensis* fighting behaviour showing typical escalation and de-escalation pathways and our classification of escalation levels.

https://www.youtube.com/watch?v=i6FA8LA2VLk

https://www.youtube.com/watch?v=fIWW4SvKvMs

**3.** *Melanotes ornata*, Orthoptera:

**<u>CLASSIFICATION</u>***: BODY SIZE ESTIMATION*; **IMPACT AND PUSH**

In the first phase (n = 39), *after a brief mutual antennal touch, individuals changed their orientation towards the opponent, subsequently elevated their body ventro-dorsally from the ground and slowly lifted their hind legs repeatedly. While doing so, opponents moved around each other gradually positioning their back towards each other while pointing the hind legs towards the opponent*. *In this first phase, opponents also accelerated antennal movements and repeatedly touched opponent's antennae and other body parts with its own antennae*. In the second phase (n = 20, Figure 1), individuals got closer**, crossed their hind legs and started a series of pushes and kick attempts**, with intermittent emission of acoustic signals. **In 3 contests, there were also frontal assaults, wrestling and biting.**

**4.** *Gryllus firmus & Acheta domesticus*, Orthoptera:

**<u>CLASSIFICATION</u>***: WEAPON DISPLAY;* **IMPACT (LEG) AND LIFT (LEG + MANDIBLE)**

When two aggressive males *meet head-on and neither retreats with the initial antennal contact, each then begins to "lash" his antennae rapidly upon the other. This is generally associated with (1) rearing the forebody, (2) drawing the palpi up and back, (3) <u>spreading the mandibles</u>, and (4) stepping forward (figs 2-3). The antennal lashing is interpreted here as the initial act of aggression in head-on encounter.*

**If neither individual retreats, the antennae are pressed together as the two crickets move forward (figs 6-9), and the two males begin "sparring" with the forelegs**. **This seems to consist of each individual striking forward with his forelegs and then jerking them back, continually placing them on top of the other male's legs and**

**hooking at his appendages with the tarsal claws** (figs 6-9). **During such sparring, either male may flip the other off-balance or butt him back with the head** (figs 11-12). The two may stand head-to-head and battle in this manner for several minutes (fig. 9). **If neither individual retreats, the mandibles are eventually locked together and there is a sort of wrestling activity (fig. IO) in which either male or both can be flipped or thrown sideways. This usually causes quick retreat, but it may also end with one male throwing the other backward up to several inches, flipping him up into the air (fig. II), or turning him completely over on his back (fig. 12)**. Sometimes both males leap into the air and come down several inches apart or some distance from their original location. Occasionally the males wrestle with locked mandibles for several seconds before one is bested. Such violent combat usually ends the fight...

**5.** *Hemideina crassidens*, Orthoptera:

<u>**CLASSIFICATION**</u>: *WEAPON DISPLAY*; **SQUEEZE AND PULL**

In 77.9% of all trials (46 of 59), **the intruder entered the gallery and extracted the resident male using his mandibles to grip the hind tibia of the resident and pull him from the cavity**. In seven trials, the resident backed out of the cavity without direct removal; these instances involved a very large resident male (head length>24 mm) and a much smaller intruder male.

*In 47.5% of trials (28 of 59), the loser either left the immediate vicinity without further contest or he antennated-palpated the opponent and then retreated (element 1).*

*Contests between male <u>H. crassidens</u> appear to occur in a single phase; fights did not proceed in an ordered manner (i.e. all behavioural elements performed in an order) with subsequences* *(different phases with different elements) of repeated displays.* Instead, **after a bout of brief and mutual antennation, males would fight (N = 31).** Fights could quickly escalate…

*Thirteen fights (22%) were settled after males faced each other and opposed their spread mandibles.*

**6.** *Rangifer tarandus*, Mammalia:

<u>**CLASSIFICATION**</u>*: WEAPON DISPLAY*; **PUSH, IMPACT AND LIFT**

1. Keep away: An individual keeps another away from a site, especially the feeding station, by its mere presence. No particular motor pattern is employed. (In the tables of this paper this behavior is included in "yielding".)

2. Supplant: Approach by one animal, followed by withdrawal or veering off by another. When focusing on the latter individual, another term for this type of interaction is "yielding."

*3. Head up: Vertical upward thrust of the head.*

*4. Head turn: Lateral twist of head that is visually amplified by the antlers ("antler dipping"). The antler tips may touch the ground. This movement seems identical with Bubenik's (1975) "head tilt."*

**5. Head lowering: Head is quickly moved vertically, often as a prelude to antler-locking (see point 1.). In the tables, this behavior is included under "antler-locking."**

**6. Pushing: Forward thrust with extended head.**

**7. Butting: Forward thrust of the forehead, often with body contact.**

*8. Lip smacking: Both sexes may rapidly open and close their mouths while the head is raised. This occurs when another individual passes closely.*

**9. Striking: Hitting with one or both raised forelegs.**

*10. Lunge: Short, quick rush at a conspecific with a sudden stop. It can be viewed as an aborted "chase." (In the tables, this behavior is included under "chase.")*

11. Chase: Driving an opponent at a fast trot or gallop over distances ranging from about 5 m to several rounds through the enclosure, totaling several hundred meters.

**12. Antler locking: Two facing animals lower their heads, touch each other's antlers, push forward and turn their heads laterally. Although dominance may be assumed for the animal remaining on the site after an antler fight, this behavior pattern was not used in the dominance matrices.**

The following three patterns typically occur during the rut:

*13. Grunt: Uttering of a short call, performed by males during the rut. It may be repeated, and results in the moving away by another male. The grunt also serves in keeping the females together in the "harem."*

*14. Tramp urinating: A rutting bull will rhythmically tramp with his hindlegs and urinate simultaneously so that urine soaks the metatarsi and hooves. In captivity, this behavior is directed at both other bulls and people.*

*15. Pawing: During the rut, the harem bull was observed to paw with his forelegs and sniff the ground when facing people. Pawing may also accompany thrashing a tree with the antlers.*

<u>Reference</u>: Müller-Schwarze, D., Källquist, L., & Mossing, T. (1979). Social behavior and chemical communication in reindeer (*Rangifer t. tarandus* L.). *Journal of Chemical Ecology*, *5*(4), 483-517.

“In contrast to the sparring that occurs at the beginning of the rut, **fights between mature males during the rut itself were often vigorous encounters in which the opponents struck with considerable force and attempted by pushing and twisting to force each other off balance**. Fights were usually terminated by one individual backing away and averting its head and antlers away from the opponent, or by both opponents stopping simultaneously.”

<u>Reference</u>: Lent, P. C. (1965). Rutting behaviour in a barren-ground caribou population. *Animal Behaviour*, *13*(2-3), 259-264.

**7.** *Capreolus capreolus*, Mammalia:

**<u>CLASSIFICATION</u>**: *WEAPON DISPLAY*; **IMPACT, PUSH AND LIFT**

*-Stiff walk: The buck lifts his feet and puts them down firmly in a demonstrative way. It is mostly performed with the side toward the other buck and from a longer distance as compared to parallel walk.*

*-Lateral display: The buck turns and shows his side to the opponent.*

*–Head shaking: The buck twists his head from left to right repeatedly, directed straight toward its opponent.*

**–Lunge: The buck jumps forward from a short distance (ca 5 m), with the antlers toward the opponent, but stops before contact, usually retreating to the original distance.**

**–Prod: The buck hits the opponent on his side or back with his antlers.**

**–Lunge/contact: The buck jumps forward from a short distance, with the antlers toward the opponent, and hits him. The opponent usually, but not always, manages to engage him with his antlers.**

**–Pushing: The bucks are head to head, trying to push each other backwards.**

**–Wrestling: The bucks lock their antlers and twist their heads from side to side, often moving around each other, possibly to make the opponent lose his balance.**

**8.** *Gallotia galloti*, Lizard:

<u>**CLASSIFICATION**</u>*: WEAPON DISPLAY*; **SQUEEZE**

*When they were close, tongue-flicking was directed at the hind part of the other's head, close to the cheek. Contests could be terminated at this stage, or even after approaching with an inflated throat by one lizard, by the other lizard fleeing. If not, 'face-off' and circling usually occurred* and then **if one of the animals did not retreat, both tried to, or bit the other and briefly rolled over, after which one of them fled and was chased by the other. Bites could be directed toward the head, trunk or base of the tail.**

**"Bite: A male opens its mouth and bites or tries to bite the opponent. The head is more frequently bitten but also the trunk or other body parts."**

Reference: Bohórquez-Alonso, M. L., Mesa-Avila, G., Suárez-Rancel, M., Font, E., & Molina-Borja, M. (2018). Predictors of contest outcome in males of two subspecies of *Gallotia galloti* (Squamata: Lacertidae). *Behavioral ecology and sociobiology*, *72*(3), 63.

**9.** *Niveoscincus microlepidotus* [*Carinascincus microlepidotus*], Lizard:

**CLASSIFICATION***: BODY SIZE ESTIMATION*; **SQUEEZE**

Male contests varied considerably in the degree of escalation and duration. *Most interactions were resolved by displays, wherein each male turned sideways towards the rival with dorso-lateral. The combat was resolved without physical contact in most of these cases.* However**, in other contests males fought viciously for up to 20 min, with severe injuries to at least one of the contestants. In several cases, males lost toes and in one instance, an entire foot was torn off**. Thus, some fights entail high costs to the interacting males.

"Basking behaviour and social interactions were recorded in all treatments over a 1 h period: preferred basking – number of minutes a lizard basked on clay basking-tile; nonpreferred basking – number of minutes a lizard basked off the clay basking-tile; **head/tail biting – presence/absence of biting;** and chasing – presence/absence of chasing."

Reference: Melville, J. (2002). Competition and character displacement in two species of scincid lizards. *Ecology Letters*, *5*(3), 386-393.

**10.** *Gonatodes albogularis*, Lizards:

**CLASSIFICATION**: *WEAPON DISPLAY*; **SQUEEZE**

*Aggressive encounters were initiated when one gecko remained still or walked slowly towards the other with a depressed throat (plus body compression); after both individuals placed with their bodies aligned laterally, they could perform 'push-ups', 'head shaking', 'whole body waving' or one of the tail displays*. This sequence could be repeated and, eventually, one of the geckos withdrew quickly from the other**. If one gecko did not withdraw after the initial displays, the other could bit and/or chase him**.

**11.** *Riptortus pedestris*, Hemiptera:

**CLASSIFICATION**: *BODY SIZE ESTIMATION*; **IMPACT, SQUEEZE AND PUSH**

In the laboratory, we observed fighting behaviors as follows. *Often, either or both males lifted their abdomens with their backs to the opponent and flapped their wings (Fig. 3A). This behavior may be a display against the opponent.* In actual fighting, **one male kicks his opponent with one or both of his hindlegs, but the kicked male shows little response (Fig. 3B); two males kick each other with their legs (Fig. 3C); one male raises his hindlegs to grasp and squeeze the opponent's body; or two males squeeze each other (Fig. 3D). The winner is the male that tries to push his opponent out of the arena and to chase him.** The loser is the male that tries to retreat from the arena. We found that no loser resumed a fight against the winner, even though observations were continued for one hour after the outcome of the contest was decided.

**12.** *Narnia fermorata*, Hemiptera:

**CLASSIFICATION**: *WEAPON DISPLAY*; **IMPACT AND SQUEEZE**

We found that males performed five distinct behaviors when paired with another male: *(1) leg display, the raising of one or both hind legs in the line of sight of another male;* **(2)**

**charge, a quick movement toward another male resulting in contact or withdrawal of the target male**; (3) mount, climbing on top of another male; **(4) kick, the use of one or both legs to quickly strike another male; (5) wrap, wrapping both legs around the body of another male.** We also recorded (6) contact, any other physical encounter not falling into these categories.

**13.** *Scopimera globosa*, Crustacea:

**CLASSIFICATION**: *WEAPON DISPLAY*; **PUSH**

Aggressive wave: One rival directed aggressive behavior to the other; *the aggressor approached the neighboring crab with its chelipeds raised upward and without extending its walking legs. This behavior was performed mainly by a burrow-holding crab towards another burrow-holding crab.*

Pre-wrestling: Both crabs were aggressive to each other, usually an interaction between a burrow-holding crab and a burrow-less crab. **The combatants faced each other and pushed the opponent with their chelipeds.**

Wrestling: Both crabs were aggressive to each other, usually an interaction between a burrow-holding crab and a burrow-less crab. Wrestling proceeded from the pre-wrestling type of aggression. **The combatants interlocked with each other, with their ventral sides touching and the first and the second ambulatory legs extended laterally as they scrambled.** Wrestling ended when one of the combatants fell on its back or retreated. **In both pre-wrestling and wrestling, the chelipeds were used for pushing or touching the opponent, but were not used to grasp the opponent.**

*14. Diogenes nitidimanus*, Crustacea:

**CLASSIFICATION**: *NO DISPLAY*; **PUSH AND SQUEEZE**

**Guarding males often extend their left major cheliped to the opponent, physically preventing rivals from approaching and sometimes flicking them away.**

*About 35% of cases (23 contests) were settled without any physical contacts when one of the males preemptively guarded the female*, while the other 64% of cases (42 contests) showed some physical contacts such as **cheliped flicking and/or grappling.**

**15.** *Austruca annulipes*, Crustacea:

<u>**CLASSIFICATION**</u>: *WEAPON DISPLAY*; **SQUEEZE AND LIFT**

Males did not grip each other with the tips of their claws; **they waited to contract their claws until they were at least partially interlaced**. Moments before they contracted their claws both males typically worked the dactyls of their walking legs deeper into the sediment. **The gripping contraction of the manus evidently was accompanied by contractions in the muscles of the carpus and merus producing a lifting force that was resisted by the opponent's leg grip into the sediment**. **If the leg grip failed the intruder sometimes was flipped upside down and thrown to the side or the resident was lifted out of the entrance of his burrow.**

**Puncture wounds on the claws we collected also showed that males forcefully grip opponents' claws when they fight.**

**16.** *Aegla longirostri*, Crustacea:

<u>**CLASSIFICATION**</u>: *BODY SIZE ESTIMATION*; **SQUEEZE AND PULL**

**-Hitting with chelipeds: An animal hits or "passes" with chelipeds on the opponent's carapace or chelipeds.**

**-Pulling/pinching with chelipeds: Consists of pressing and pulling quickly the cheliped (or the pereiopods) of the opponent. The animal tries to catch the opponent, but it cannot, during this act the antennae (generally) remain horizontal to the body.**

**-Holding (catching) with chelipeds: Consists of holding (catching) the opponent with chelipeds. Generally the part which is held is the chelipeds of the other animal or even the pereiopods and antennae, and sometimes the animals try to catch the opponent's cephalothorax.**

*-Whipping with antennae: This is a quick below with the antennae toward the back of the body, and the opponent is behind the performer of the act.*

*-Touch with antennae: Consists in touching the opponent quickly with the antennae; the opponent being near the front of the animal, the touch occurs both on the body and the antennae of the opponent (commonly observed during combat).*

**-Pushing the opponent/displacing the body: One animal tries to displace the other pushing it with its own body (abdomen or cheliped).**

**-Turning the opponent upside down: During the combat an animal turns the opponent over, leaving it with the ventral part of the body up. (VERY RARE BEHAVIOR)**

-Going up the opponent: An animal goes totally up over the opponent's body, even when one of them is climbing the arena.

-Chasing: An animal chases (quickly approaching) the opponent, while the latter attempts to escape.

Reference: Ayres-Peres, L., Araújo, P. B., & Santos, S. (2011). Description of the agonistic behavior of *Aegla longirostri* (Decapoda: Aeglidae). *Journal of Crustacean Biology*, *31*(3), 379-388.

**17.** *Cherax dispar*, Crustacea:

**CLASSIFICATION**: *WEAPON DISPLAY*; **SQUEEZE AND PUSH**

Male crayfish routinely use their enlarged front claws (chelae) for both intimidation and fighting, making these organisms ideal for studying the honesty of aggressive signals (fig. 1). *During competitive interactions, we frequently observed males displaying their chelae to opponents, but direct physical contact occurred only in a small proportion of interactions.*

Fights were defined as two animals facing each other with each attempting to hold and unbalance the other. **Crayfish typically used their chelae to push their opponent and to take hold of the adversary's chelae**. Eventually, one of the contestants would move away, and the remaining animal was scored as the winner of the fight.

**18.** *Dicronocephalus wallichii*, Coleoptera:

**<u>CLASSIFICATION</u>***: BODY SIZE ESTIMATION*; **LIFT AND PUSH**

The owner abruptly turned to an approaching intruder ('Sense orientation'), even when the intruder came from behind. *The owner stretched and touched the intruder's body or forelegs and subsequently intensely moved his forelegs ('Tapping', Fig. 2A, F). The intruder also showed tapping with the forelegs in response to the owner's behaviour. After a few seconds of tapping, the intruder walked away from the owner in many cases ('Intruder escapes', Fig. 2F).* If the intruder continued to approach and/or tried to mount the female, **the owner pushed him away with his horns ('Push') or pried him away from the female's back or substrate with his horns ('Pry',** Fig. 2B, F and Movies 1, 2). *The forelegs of the owner moved intensely and touched the opponent's body during 'Push' and 'Pry'.* The contests usually began with 'Tapping', but this step was occasionally skipped, especially when the intruders directly flew to the guarding males or dashed into them (Fig. 2F). **During 'Pry', the intruder resisted the owner's movements, and the two males tussled for a few seconds to one minute, holding each other's horns tightly** (Fig. 2B, F). **The intruder escaped ('Intruder escape', Movie 2) or was flipped away ('Flip', Movie 3) in the end**. While the owner lifted the intruder's body to flip him, the intruder struggled to cling onto the owner's body. The owner rapidly moved his forelegs and midlegs to disengage the intruder's body from himself (Fig. 2C). The owner was found to drive the intruder c. 10–50 cm away from his mate using the horns and forelegs during 'Push' or 'Pry' (Movie 3). In such cases, the owner returned to his mate immediately after winning the fight. Females frequently (49 out of 140 cases that escalated into 'Pry' or 'Push') tried to escape from the owner during the prolonged fighting (Table 2).

The owner was flipped away by the intruder, and takeover occurred ('Owner flipped') after 'Pry' (Fig. 2F). In some cases, the owner and intruder were found to fall together during escalated tussling ('Fall with intruder') (Fig. 2F). The owner accidentally fell together with his mate during the contests in other cases ('Fall with female') (Fig. 2F). **The males were never obviously injured during Fighting.**

**19.** *Trypoxylus dichotomus*, Coleoptera:

<u>**CLASSIFICATION**</u>*: NO DISPLAY*; **LIFT AND PUSH**

When two individuals on the same feeding spot came in contact with a body part of each other, it was defined as 'Encounter' (stage 1). **Immediately after two males encountered, they always faced and shoved each other with their horns (stage 2: 'Shoving'**). Then, in 72 of the 124 cases, one male began to retreat and was chased and thrust with a horn by the other male, and finally the former male was driven away from the feeding spot (stage 4A: ' Chasing' ). In the other 52 cases, **after 'Shoving', males put their horns under the opponents' bodies each other and pushed stronger trying to lift and flip up the opponent by flexing their horns dorsally (stage 3: ' Pry'** ). In 24 of the 52 cases, **one male was immediately flipped up from the feeding spot and the interaction was terminated quickly** (stage 4B: 'Flipping'). In the other 28 cases, interactions were not terminated so quickly. Two males pushed each other intensely keeping their postures as in stage 3 for a while. After a while, one male suddenly began to retreat, and the interaction proceeded to stage 4A and was terminated.

**20.** *Heterochelus chiragricus*, Coleoptera:

**<u>CLASSIFICATION</u>**: *WEAPON DISPLAY*; **SQUEEZE AND LIFT**

*Sometimes, a mate-guarding or in copula male would lift his hind legs and vibrate his feathery tarsi at the appearance of an intruding male* (Figure 1). **Facing away from each other, opponents lashed out with their hind legs, reflexively clenching their tarsi in a lever-like action, attempting to dislodge their opponent either off** a female and/or out of the flower arena. Resident males would cling to females using their fore legs and midlegs while battling with their hind legs. Fight duration ranged from 2 to 120 s (mean duration ± SE = 18.68 s ± 1.68). **Contests were generally won when one contestant used his hind legs to grasp his opponent and lever him off the female and out of the flower**. In some instances, intruders simply gave up and left the flower. **Males utilized their enlarged tibia and femurs in conjunction as levers when lifting opponents.**

**21.** *Sagra femorata*, Coleoptera:

**<u>CLASSIFICATION</u>**: *NO DISPLAY*; **SQUEEZE AND LIFT**

**During combat, males attack one another, using their hindlegs to squeeze rival males, pry apart copulating pairs, and steal mates**.

*Interactions began as males aggressively approached one another, progressed to one of five "escalation levels"*, and ended when one contestant either retreated or was forcibly removed from the fighting area. Escalation levels were defined as follows: Level 1) non-violent interaction, Level 2) violent interaction without full combat, Level 3) full combat of mild intensity, Level 4) full combat of high intensity where one contestant retreats, Level 5) full combat of high intensity where one contestant is forcibly removed from the fighting area.

*When a male (intruder) encountered a male (resident) and female pair, he attempted to mount the mating pair to disturb their copulation and to mate with the female* (Fig. 3b). **However, the resident male raised his hind legs to the intruder. In two of the five observations, the resident male succeeded in guarding the female and continued mating, while the intruder retreated to search for another mate**. In contrast, three resident males allowed the intruders to mount their own backs, at which point the intruder grasped the resident male by the hind legs and swung the resident bodily off the female (Fig. 3c, d). Of these three, two resident males failed to guard the females, the genitalia coupling between the resident male and the female was interrupted and the males retreated from the females (Fig. 3d-f). One resident male ultimately succeeded in protecting his mate from the intruder.

Reference: Katsuki, M., Yokoi, T., Funakoshi, K., & Oota, N. (2014). Enlarged hind legs and sexual behavior with male-male interaction in *Sagra femorata* (Coleoptera: Chrysomelidae). *Entomological news*, *124*(3), 211-220.

**22.** *Aegus chelifer*, Coleoptera:

**CLASSIFICATION**: *BODY SIZE ESTIMATION*; **LIFT, PUSH AND SQUEEZE**

*Aggression between males always occurred after physical contact that involved touching their antennae onto the opponent's body, and could be classified into the four levels of aggressive intensity* (Table 2) (level 0: no aggression, level 1: aggressive posture, level 2: one sided-attack and level 3: wrestling). From a total of 191 tested pairs, 118 male pairs were found to display aggressive behavior (level 1, 2 or 3), with 18, 20 and 80 pairs showing a maximum level of aggression of level 1, 2 and 3, respectively (Online Resource 3). **The main fighting style was prying and lifting the opponents from the ground using mandibles**. **Other fighting styles were also observed during the combat, such as pushing and biting.** While wrestling, both males might stop their fight for several seconds and then re-engage again (videoclip at Online Resource 4). From a total of 100 contests with fights at an aggression level of 2 or 3, the outcome of the fight could not be defined for six contests because both males stopped the fights simultaneously and then retreated away from each other. **We observed visible injuries in one of the males in two contests, where one male lost a front leg tarsus and another male lost a hind leg tarsus. However, note that this was because their legs tightly grabbed the floor when the opponents were trying to pry and lift them, and was not caused by directly by the opponent.**

**23.** *Ficedula hypoleuca*, Aves:

**<u>CLASSIFICATION</u>***: BODY SIZE ESTIMATION*; **SQUEEZE AND PIERCING**

However, in two conflicts (cases 8 and 9; Table 1) **the females tumbled around on the ground, trying to hold each other with their legs and to attack each other by pecking,** for a total amount of time of 12 and 25 min, respectively.

In summary, *conflicts may be generalized as starting with physical fighting after which the female that turned out to be the winner defended the nestbox*, *whereas the loser after some time often started intense bouts of alarm calling*. However, in several conflicts there were intermittent periods in which the behaviour of the females could not be seen or heard on the videotapes. *Field observations of conflicts suggest that these periods often involved females sitting still and watching each other, often at very close range and sometimes in threat postures.*

**24.** *Cambridgea foliata*, Arachnida:

<u>**CLASSIFICATION**</u>*: BODY SIZE ESTIMATION;* **PUSH**

When beginning their interactions, *if the spiders were far apart, one male would pull at the web with all legs at once, creating a large distortion in the web ('shaking' the web). If the spiders were closer to each other, they drummed their palps or their first pair of legs on the web*. If this behaviour was only performed by one spider, we described this as a 'one-sided' interaction and categorized it as intensity 1. When both rivals engaged in this behaviour, it was defined as 'signalling' and categorized as intensity 2. *We called all noncontact communication 'signalling' rather than displaying to emphasize that the signals are strictly nonvisual.*

Males **entering a sparring phase reared up on their hind pairs of legs and pushed at each other with their anterior pairs of legs** (Fig. 2a). We recorded the first male to enter this posture as the 'initiator' of contact. This phase could last for several seconds to a few minutes. Occasionally, **fights escalated to grappling whereby males would place all legs back on the web, lock chelicerae and push at each other** (Fig. 2b)

"The contest behaviours observed in *C. plagiata* followed a clear pattern of escalation. Initially, *males oriented towards each other, hanging beneath the web, and signalled with bouts of stridulation (using a peg on the ventral side of the pedicel against ridges on the abdomen). This progressed to males approaching each other and tapping the rival cepholathorax with their* *forelegs.* Following this, **males showed the highest intensity behaviour recorded (grappling) by opening their chelicerae widely and locking them with those of their opponent. During grappling, males also locked legs with the opponent. Contests were resolved once a male retreated and turned to flee the web.** The winning male often gave chase and, in rare cases, stridulated."

Reference: McCambridge, J. E., Painting, C. J., Walker, L. A., & Holwell, G. I. (2019). Weapon allometry and phenotypic correlation in the New Zealand sheetweb spider *Cambridgea plagiata*. *Biological Journal of the Linnean Society*, *126*(2), 349-359.

**25.** *Tetranychus urticae*, Arachnida:

**CLASSIFICATION**: *WEAPON DISPLAY*; **PUSHING AND PIERCING**

The agonistic behavior exhibited by male *U. formosa* in some respects resembles that of the spider mite *Tetranychus urticae* Koch, males of which vigorously guard quiescent female deutonymphs (Potter 1981).

Reference: Dimock Jr, R. V. (1983). In defense of the harem: intraspecific aggression by male water mites (Acari: Unionicolidae). *Annals of the Entomological Society of America*, *76*(3), 463-465.

*A fight may be initiated by either individual by confronting the other with outspread front tarsi. The challenged male either backs away or responds by raising and spreading apart his forelegs and extruding his cheliceral stylets. The combatants circle* and **rush each other, flailing their forelegs and jousting with their extruded stylets. There is much pushing and**

**grappling.** Mites often use their palpal glands to apply strands of silk to the mouth parts and legs of the opponent. An individual may be so badly entangled that his movements are impeded and he retreats to clean himself, thus ending the encounter, at least temporarily. Rarely, **one male punctures the other's integument with his stylets. Males injured in this way are crippled and death usually results. Although very few agonistic encounters of the fourth type were observed in progress, dead deflated males were not unusual in the vicinity of guarded females.**

Reference: Potter, D. A., Wrensch, D. L., & Johnston, D. E. (1976). Aggression and mating success in male spider mites. *Science*, *193*(4248), 160-161.

**26.** *Kosciuscola tristes*, Orthoptera:

CLASSIFICATION*: WEAPON DISPLAY*; **IMPACT (LEG) AND SQUEEZE (MANDIBLE)**

During fights, males displayed several distinct behaviours: **bite (mandibles engage with another grasshopper's body), kick (hindlegs in a sharp movement away from the body resulting in the propulsion of another)**, *mandible flare (grasshopper arches back shakes head and opens mandibles),* **mount (grasshopper jumps from within 10 cm and lands along the anterio-posterior axis of the dorsum of another) and grapple (grasshoppers lock legs and roll around)** (Figure 1, Table 1).

**During fights, we observed bites, kicks, mounts and mandible flares** (the latter only executed by the defender) (see Movie S1). Challengers were frequently seen mounting other challengers and mounting the defender (Table 1). **Defenders and challengers often exchanged bites (Figure 1a, Table 1), which in some cases caused immediate visible damage.**

*Defenders frequently reared back and flared their mandibles at challengers, but challengers never flared their mandibles at defenders* (Figure 1b Table 1). *During mandible flaring the defending male arched back and shook his head while expanding his white maxilla and labrum to expose his black mandibles and his mouth* (see Movie S1).

Challengers often mounted defenders, who never responded in a like manner, as doing so would relinquish their position on top of the female (Table 1).

**27.** *Carcinus maenas*, Crustacea:

<u>**CLASSIFICATION**</u>: *WEAPON DISPLAY*; **IMPACT, SQUEEZE AND PUSH**

CHELIPEDS IN, BODY RAISED - The body of the crab is raised as high as possible over the substratum by the fully extended legs. *The chelipeds are folded in front of the cephalothorax or occasionally pointed downwards.*

*CHELIPED DISPLAY - The body of the crab is raised as high as possible over the substratum using the fully extended legs. Chelipeds are held out in front with the chelae open or closed.*

*LATERAL MERUS DISPLAY - As cheliped display but the chelipeds are held out at 180 degrees to the body with the chelae open.*

**STRIKE - One crab suddenly hits out at the other with one or both chelipeds.**

**GRASP - When one crab uses its chelae to pinch the carapace, chelipeds or legs of the other crab.**

CLIMB ON - One crab attempts to or actually climbs on top of the other crab. When on top of the opponent the chelae are usually directed in front of the cephalothorax of its opponent and grasping may occur.

CLIMB OFF - After successfully climbing on, the crab climbs off the other crab.

**PUSH - The crab pushes its opponent using its chelipeds, pushing forward with the walking legs and rubs the body of its opponent or grasps using the chelae; or pushes the other crab away using the chelipeds only.**

Reference: Sneddon, L. U., Huntingford, F. A., & Taylor, A. C. (1997). The influence of resource value on the agonistic behaviour of the shore crab, *Carcinus maenas* (L.). *Marine & Freshwater Behaviour & Phy*, *30*(4), 225-237.

**28.** *Hipposideros armiger*, Mammalia:

**CLASSIFICATION**: *BODY SIZE ESTIMATION*; **IMPACT**

*-Ear movements: Ears move around to detect the opponent; often accompanied by echolocation. There is no physical contact.*

*-Head raise/turn: Contestants raise and/or turn heads by bending their neck backward toward the interactive bat. There is no physical contact.*

*-Crouching, unfolding wings: Opponents perform an upside-down crouch, stretch necks and initiate rapid wing flapping toward the interactive bat. Sometimes one of them walks toward the other. Often one or both contestants start to emit social calls. There is no physical contact between opponents although sometimes the distance between contestants is less a wing length.*

**-Boxing: Boxing is an escalating and physical fight between two males. Each of them uses the carpal joint to knock each other vigorously until one of them retreats.**

**-Wrestling: Bats briefly holding on to each other with either their forearms or semi-extended wings.**

**29.** *Cambarus robustus*, *Cambarus carinirostris*, *Faxonius obscurus*, Crustacea:

<u>**CLASSIFICATION**</u>: *BODY SIZE ESTIMATION*; **SQUEEZE AND PUSH**

Formal descriptions were provided by Zackary Graham, the first and corresponding author of the paper that analysed the fighting behaviour of these three species (and is referenced below).

"These crayfish start the contest with antennal whipping like most crayfish, which is followed by claw interlocking. During claw interlocking, crayfish push each other using their claws. When fights get more intense, crayfish start pinching each other and squeezing the rivals until one of them withdraws. Injuries were not observed."

<u>Reference</u>: Graham, Z. A., & Angilletta, M. A. (2020). Separating noise and function in systems of animal communication: a comparative study of aggressive signaling in crayfish. *bioRxiv*. DOI: https://doi.org/10.1101/2020.08.03.234419

**30.** *Librodor japonicus*, Coleoptera:

<u>**CLASSIFICATION**</u>: *DISPLAY WEAPON*; **SQUEEZE, PUSH, PULL AND LIFT**

Male interactions usually occurred on the banana fruit or on the sap site and were classified into four levels as follows: level 1 (no reaction), where neither male acted against the adversary even if one male contact with the other, where one or both males hid under the banana fruit, the sap site, or the filter paper; *level 2 (warning display), where one male responded by moving its mandibles or its head up and down facing the opponent - here, the head movement frequently made by the resident male toward the intruder regardless of the opponent's sex might be considered a response to broad external stimuli* *including approach of the females or the other insects - the warning stance was seldom observed in females*; level 3 (one-sided attack), where one male responded to the opponent by aggressive **behavior, including spreading the mandibles and mounting the opponent - at this stage, the attacking male bit or pushed his opponent with his mandibles, but the attacked male showed little response; and level 4 (escalated fighting), where two males came into bodily contact and attacked each other**. **Escalated fighting was further classified into three types: (i) males faced each other, interlocked their mandibles and shoved each other; (ii) males repeatedly grabbed a hold of each other with the mandibles and pulled, trying to lift the rival off the substrate; and (iii) males opened their mandibles and bit each other**. The outcome of the contest was decided only at levels 3 and 4. The winner was considered the male that pushed his opponent out of the fighting site and chased him. The winners chased rivals off the banana slices or the sap sites and pursued them. The loser was the male that retreated from the fighting site.

**31.** *Gnatocerus cornutus*, Coleoptera:

<u>**CLASSIFICATION**</u>: *WEAPON DISPLAY*; **SQUEEZE, PUSH AND LIFT**

In the laboratory, males of *G. cornutus* frequently fought each other with their mandibles. Male interactions were classified into two levels as follows: 1) level 1 (one-sided attack), where one male responded to the opponent with aggressive behavior. **The attacking male bit or pushed his opponent with his mandibles**, but the attacked male showed little response, and 2) level 2 (combat), where two males came into bodily contact and attacked each other. Combat was further classified into three types: (i) **males faced each other, interlocked their mandibles and shoved each other** (Fig. 3a); **(ii) males repeatedly put their mandibles under the opponent's body, trying to lift**

**the rival off the substrate (Fig. 3b), and (iii) males bit and shoved each other with their mandibles (Fig. 3c).**

*Two males often faced each other and moved their heads with enlarged genae up and down. This behaviour may be display against the opponent*. Alternatively, two males could come into bodily contact and attack each other. This fighting behaviour was further classified into three types: (1) **males faced each other, interlocked their mandibles, and shoved each other; (2) males repeatedly put their mandibles under the opponent's body, trying to lift the rival off the filter paper; and (3) males bit and shoved each other with their mandibles.**

Reference: Okada, K., & Miyatake, T. (2009). Genetic correlations between weapons, body shape and fighting behaviour in the horned beetle *Gnatocerus cornutus*. Animal Behaviour, 77(5), 1057-1065.

**32.** *Parastacus brasiliensis* and *Parastacus pilimanus*, Crustacea:

**CLASSIFICATION**: *BODY SIZE ESTIMATION;* **PUSH, PULL AND SQUEEZE**

Ethogram codes for agonistic behaviour in *Parastacus*. Format, score given: behavior.

-2: Retreat with a tail flip.

-1: Retreat by walking away from the opponent.

0: Ignore the opponent/non-aggressive behaviours.

*1: Approach without agonistic display.*

*2: Approach with meral spread and/or antennal whip.*

**3: Aggression with closed chelae: touching, punching and pushing the oponent.**

**4: Active use of the chelae to grab the opponent's appendages, or chela strike.**

**5: Intense combat: animals performing several agonistic acts simultaneously, trying to grab and pull the opponent's body parts, or attempting to turn/carry the opponent.**

**33.** *Gryllus pensilvanicus*, Orthoptera:

<u>**CLASSIFICATION**</u>: *BODY SIZE ESTIMATION*, **PUSH AND SQUEEZE**

Fighting crickets display a stereotyped sequence of escalating motor behaviours. We divided this escalation cascade into the following levels of aggression (modified after Alexander 1961; Fig. 1): Level 0: mutual avoidance, no interaction; Level 1: clear dominance, one animal retreats immediately*; Level 2: antennal fencing, involving both animals; Level 3: mandible spreading (unilateral), one animal displays spread mandibles*; *Level 4: mandible spreading (bilateral), both animals display spread mandibles;* **Level 5: mandible engagement, mandibles make contact and crickets push against each other; Level 6: 'wrestling', an all-out fight where the animals interlock mandibles and push each other. They may repeatedly disengage, struggle for position, bite other body parts, and re-engage mandibles to push the opponent.** The fight can be concluded at any of the levels 1–6 by one opponent, the loser, retreating, upon which the established winner typically produces victory displays such as the rivalry song and characteristic body-jerking movements.

<u>Reference</u>: Hofmann, H. A., & Schildberger, K. (2001). Assessment of strength and willingness to fight during aggressive encounters in crickets. *Animal Behaviour*, *62*(2), 337-348.

**34.** *Anolis*, Squamata:

**<u>CLASSIFICATION</u>**: *WEAPON DISPLAY*, **SQUEEZE**

*The 'dewlap display' consisted of one or more consecutive bouts of the lizard extending its dewlap, with each extension usually accompanied by a cluster of rapid up and down movements of the front of the body (bobbing). A single bout consisted of one extension and retraction of the dewlap. The 'bobbing display' consisted of bouts of bobbing without the extension of the dewlap. A single bout consisted of several rapid up and down movements, with pauses of more than 0.5 s separating bouts.* **'Sparring' consisted of a lizard opening its jaws or biting its opponent. Usually, this involved each lizard grappling for a grip on its opponent's jaws, but often involved into a situation where each lizard held its opponent by the jaws for an extended period of time.** A dewlap display or bobbing display began with the first bout of that display and ended with the first bout of another aggressive behaviour, or when the contest was ended. For sparring, only one bout was counted between initiation of sparring and the initiation of another behaviour.

Reference: McMann, S. (1993). Contextual signalling and the structure of dyadic encounters in *Anolis carolinensis*. *Animal Behaviour*, *46*(4), 657-668.

**35.** *Ctenophorus maculosus*, Squamata:

**<u>CLASSIFICATION</u>**: *WEAPON DISPLAY*, **SQUEEZE**

Behaviour: Description

Approach: Moves slowly towards opponent;

**Bite: Bites any body part on opponent;**

Chase: Runs towards and pursues opponent

*Head bob: Pronounced up and down 'nodding' movement of the head*

*Push-up: Lateral compression of the body and a raised stance*

**Tail lash: Tail lashed at opponent**

**36.** *Onthophagus taurus* and *Onthophagus acuminatus*, Coleoptera:

**CLASSIFICATION**: *WEAPON DISPLAY*, **PUSH AND LIFT**

*After initial contact between contestants, both the defending resident and the intruding male assumed a typical fighting position with the head and the thorax held low, the abdomen held high, and the legs braced against the tunnel walls* (Fig. 2). *This lowering of the head resulted in head horns pointing towards the opponent*. Horns in *O. taurus* consist of two long, bow-shaped structures (Fig. 1), **and males engaged in head-to-head contact tightly embraced the thorax of their opponent with their horns** (19/19 competitions, Fig. 2). **At this stage both males vigorously pushed each other while performing frequent and rapid upward jerks with their heads** (19/19 competitions). **Clicking sounds could be heard clearly through the glass panes as the head and horns of both males came into contact during head-to-head combat. Fights continued in this fashion until one male was able to dislodge his opponent sufficiently from the substrate. Once this was achieved, subsequent forward pushes and upward jerks allowed the stronger male either to drive his opponent out of the tunnel (if the stronger male**

**started out lower in the tunnel) or to push his opponent further into the tunnel until the tunnel diameter permitted the stronger male to climb around his opponent and then force the opponent out of the tunnel.** In all cases, fights ended when one male left the tunnel.

**37.** *Neogonodactylus bredini*, Hoplocarida:

**<u>CLASSIFICATION</u>**: *WEAPON DISPLAY*, **IMPACT**

*Some mantis shrimp species use a visual display* and **the ritualized exchange of strikes** during territorial conflicts (reviewed in [10]). *During the 'meral spread' visual display, the raptorial appendages are spread laterally and ventrally such that several parts of the appendage are presented to the competitor and the individual displaying the meral spread is biomechanically unable to strike* (figure 1) [10]. *The meral spread is considered a signal of aggressive motivation [10–12] and possibly performance* [1]**. In addition, competitors exchange strikes using a ritualized 'telson coil' behaviour [10,13], in which the receiver of a strike coils its tailplate, or telson, in front of its body to receive the blow.**

**38.** *Teleogryllus commodus*, Orthoptera:

Same description for *Acheta domesticus*

**39.** *Anisolabis maritima*, Dermaptera:

**<u>CLASSIFICATION</u>**: *WEAPON DISPLAY*; **IMPACT AND SQUEEZE**

"High-intensity interactions often occurred immediately following first contact. In open sites (petri dishes), *antennal contact* was followed by **turning and striking or by curving the abdomen forward and then striking; only occasionally did a male strike directly backward.** In contrast, in tunnels, where the animals were less mobile, males struck by first turning 180° and then moving directly backward. **Pinches were frequent in both situations**. They lasted longer in tunnels than in the open. **Pinches in the open usually occurred on the posterior portion of the abdomen (Figure 57), whereas those in tunnels occurred often on both the abdomen (Figure 58) and the thorax**. Simultaneous pinches were rare. **In three cases a male in the open lifted and shook a pinched opponent; in one case he then threw him against the wall of the dish (Figure 5). In some cases one male struck another with a lateral or a backward slam and knocked him some distance** (Figures 59, 60). When pinched, the opponent struggled to free himself by pulling, twisting his abdomen, returning the pinch, or biting the other male. Another tactic with which males freed themselves on several occasions was for the pinched male to turn and move forward under the abdomen of the other male, thus twisting the pinching male's abdomen sharply (Figure 61).

*Low-intensity battles both in the open and in tunnels included presentations and short pushes with the cerci. Opponents were struck with the tip of the abdomen with the cerci spread. Winners of interactions in the open often swung their abdomens strongly from side to side after the opponent withdrew.* Losers in tunnels were less likely to withdraw, rather they simply remained more or less immobile with their cerci toward the opponent."
